# Identifying the non-Bifidobacterial hallmarks of a Bifidobacterium-receptive gut microbiome

**DOI:** 10.1101/2025.09.09.675069

**Authors:** Sourav Goswami, Alisha Ansari, Chetan Saraf, Paul W. O’Toole, Fergus Shanahan, Vineet Ahuja, Tarini Shankar Ghosh

**Author notes:** Equally contributing authors (Sourav Goswami, Alisha Ansari).

## Abstract

Bifidobacteria are key health-associated members of the human gut and widely used as probiotics, but their colonization success varies across individuals, partly due to baseline microbiome composition.

In this analysis encompassing 51,058 gut microbiomes (148 cohorts spanning 46 countries across six continents), we investigated non-Bifidobacterial features associated with the overall detection and abundance of eight major Bifidobacterial species. While non-Bifidobacterial community composition significantly correlated with Bifidobacterial detection in 74% of analyzed cohorts, the associations for the individual species showed distinct age-/lifestyle-/disease-specific patterns (greater similarity within similar cohorts). We quantified these relationships as Association-Scores, further stratified by age group, lifestyle, profiling strategy (16S/WGS) and nine diseases, and reasonably reproduced in an external validation-cohort-set of 8 *Bifidobacterium* intervention studies (1,063 gut microbiomes).

We subsequently utilized these scores to build microbiome-level receptive-scores corresponding to the 8 major Bifidobacterium species and their overall detection. Both robust linear regression and cross-study validation showed that combining baseline abundance of the administered *Bifidobacterium* with the corresponding baseline receptive-scores significantly predicted post-treatment persistence/increase in 75% trial-probiotic pairs. We identify 129 species-specific-genome-level functional-groups in non-Bifidobacterial taxa correlating with Bifidobacterial Association-Scores.

This global analysis highlights non-Bifidobacterial microbiome features influencing Bifidobacterial colonization and provides tools to predict personalized responses to Bifidobacterial-probiotic interventions.

## Introduction

Bifidobacteria are one of the earliest colonizers of the human gut where they feed on human milk oligo-saccharides and exhibit health-associated activities^1^. Benefits include the promotion of pathogen resistance, immune maturation, anti-inflammation, and microbiome maturation including the promotion of butyrate producers^2^. After infancy, Bifidobacteria have also been associated with beneficial influences including: enhanced nutrient absorption; immune regulation; protection against infection, inflammation and cancer; bone-health; cardio-metabolic health and fat metabolism, and psychological welfare^3,4^. Consequently, various species belonging to the genus *Bifidobacterium* have been explored as potential probiotics (both independently or as part of dietary supplements and functional foods; as single microbe or in consortia)^3,5^. According to definition by International Scientific Association for Probiotics and Prebiotics (ISAPP), probiotics are “live microorganisms which when administered in adequate amounts confer a health benefit on the host”^6^. *Bifidobacterium* species such as *B. animalis*, *B. longum*, *B. breve*, *B. bifidum* and *B. adolescentis*, have been assessed for probiotic efficacy in a diversity of disorders^7–9^. However, a continuing challenge is the inconsistency and variability of *Bifidobacterium*-derived probiotic products in terms of efficacy and the extent of colonization of the administered *Bifidobacterium* taxa^10,11^. To rationally select *Bifidobacterium* strains for future trials, systematic reviews and cross cohort data analysis of endogenous *Bifidobacterium* species across different age-groups and life-style patterns have been conducted^12–14^. These reveal variability in *Bifidobacterium* taxa across different age-groups and life-style patterns (especially industrialization). For example, *B. breve* and *B. bifidum* are predominant in the infant microbiome, whereas *B. adolescentis* is typically more prevalent in the adult gut, and *B. longum* is present in all age-groups but with specific strains associated with infants and adults^13^. In general, Bifidobacterial species are enriched in the westernized populations when compared with non-westernized, although the relative abundances of *Bifidobacterium* are similar in infancy^12^. Overall, a lower *Bifidobacterium* abundance has been found in elderly subjects. Other than these general associations, research trials have shown that persistence of individual *Bifidobacterium* strains varies widely even within individuals of a similar age-group and life-style patterns^10,11,15^. This variability has been attributed to characteristics of the host’s baseline gut microbiome.

A more comprehensive understanding of the degree to which the baseline gut microbiome (particularly non-Bifidobacterial components) influences the representation of different *Bifidobacterium* members is needed. Large-scale data-driven studies are required to systematically investigate how individual non-Bifidobacterial members of the baseline gut microbiome associate with different *Bifidobacterium* taxa in hosts of different age-groups and life-styles and if genome-specific functions influence such association-patterns? Moreover, leveraging such data to stratify individuals at baseline according to their likelihood of responding to different probiotic *Bifidobacterium* treatments is an intriguing possibility.

To address these issues, we systematically investigated 51,058 gut microbiomes across 46 countries to investigate the non-Bifidobacterial microbial members associated with the overall detection and abundance of eight major Bifidobacterial taxa, across 148 cohorts of different age-groups, industrialization-patterns, disease-groups and microbiome-profiling strategies. The non-Bifidobacterial community composition had a strong influence on the Bifidobacterial composition of the gut microbiome, with the association-patterns of the individual non-Bifidobacterial taxa with the different Bifidobacterial species showing distinct patterns depending upon study-cohort characteristics. We quantified these association-patterns as Association-Scores, further stratified by age group, lifestyle, disease and profiling strategy (Amplicon/16S and Whole Genome Sequencing/WGS). These Association-Scores were finally utilized to formulate microbiome-level Receptive-Scores for the 8 major *Bifidobacteria* and their overall detection. Combining baseline abundance of the administered *Bifidobacterium* along with the corresponding baseline Receptive-Score significantly predicted post-treatment persistence/increase across 82% trial-probiotic pairs from the 8 additional intervention-studies (encompassing 1032 gut microbiomes).

## Results

### Global Bifidobacterial prevalence patterns are strongly associated with study-cohort characteristics and the composition of non-Bifidobacterial microbiome

We initiated the investigation by building a global collection of 45,809 gut microbiomes collected from 121 study-cohorts (a subset of the global collection of 51,058 gut microbiomes), spread across 46 countries (spanning all six continents) (**Methods**; **Table S1**; **Figure S1A**; also described in ^16^). We specifically focused on gut microbiomes from subjects in three distinct age-groups, namely ‘Infant/New-Born’ (Age <= 2y, n=9,123 gut microbiomes), ‘Adult’ (18y <= Age < 60y; n=26701 gut microbiomes) and ‘Seniors’ (Age >= 60y, n=9985 gut microbiomes). For every cohort, we also collected the information pertaining to its cohort-type (‘IndustrializedUrban’, ‘Mixed’ and ‘RuralTribal’), as described earlier^16,17^.

We identified 42 Bifidobacterial taxa that were detected in at least one microbiome. Amongst these, eight, namely *B. longum*, *B. adolescentis*, *B. pseudocatenulatum*, *B. bifidum*, *B. dentium*, *B. breve*, *B. catenulatum* and *B. animalis* demonstrated strong associations with the gut microbiomes with prevalence rates of >=25% in >= 5% of studies (in at least one age-group), in line with previously reported observations^12–14^ (**Figure S1B**). Both age-group and life-style (especially Westernization/Industrialization versus Traditional life-style) have significant impact on the gut microbiome, which were also observed in our investigation^18,19^. We replicated several previously observed trends pertaining to the prevalence of these taxa. For example, there was significant variation in the prevalence patterns of the different Bifidobacterial species between the Infant and Adults/Senior Cohorts (PERMANOVA R-squared=0.09, P=0.001) (**Figure S1C**). *B. breve/bifidum* were especially prevalent in infant cohorts and *B. adolescentis/animalis/catenulatum* were more prevalent in the adult/seniors (as reported earlier^13^) (**Figure 1**; **Figure S1B-D**). The prevalence patterns also showed significant variations across studies belonging to the cohort-types (PERMANOVA R-squared=0.05, P=0.05, **Figure S1E**). In general, there was an overall reduced prevalence of Bifidobacterial taxa in the RuralTribal as compared to Westernized (IndustrializedUrban) cohorts (as observed earlier^12^) (**Figure S1D-E**).

**Figure 1.**
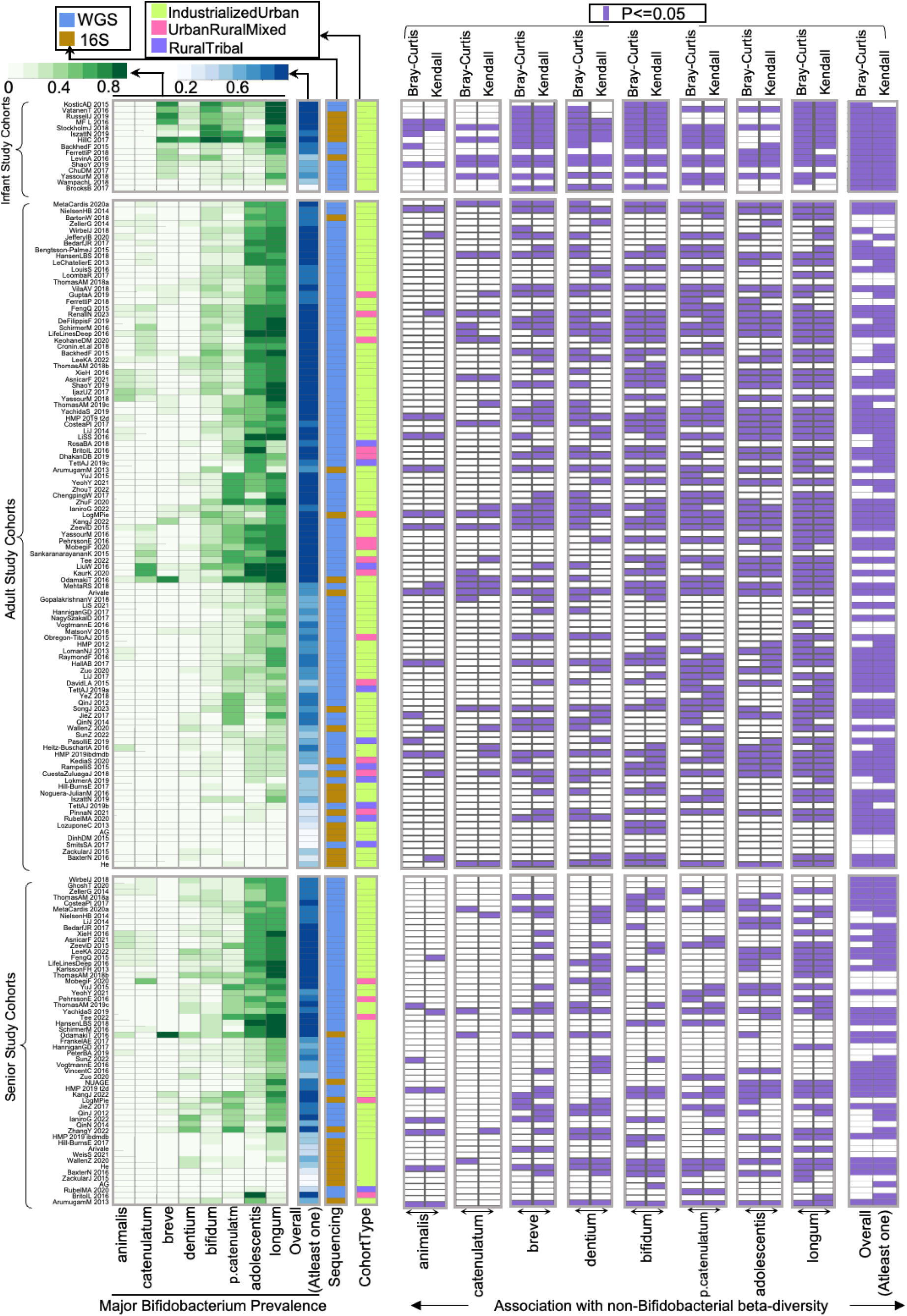
Prevalence patterns of Bifidobacterial taxa across the investigated study-cohorts, along with the study-specific associations of the non-Bifidobacterial gut microbiome with the representation of the major Bifidobacterial species. (Left Panel) Heatmap showing the prevalence patterns of the eight major Bifidobacterial species and the fraction of microbiomes with at least one Bifidobacterial species detected (with abundance >= 0.0001) across the different study-cohorts. The study-cohorts have been divided based on the age-groups (Infant, Adult, Senior). Also indicated are the sequencing-types (WGS or 16S) and the cohort-type (IndustrializedUrban, UrbanRuralMixed, RuralTribal). Panels on the right indicate the associations of the non-Bifidobacterial community compositions with the representations of the eight different Bifidobacterial species as well as the overall *Bifidobacterium* detection rate. The associations were computed in each study-cohort using PERMANOVA with the non-Bifidobacterial community variations using two different distance measures, namely Kendall and Bray-Curtis. The eight major Bifidobacterial species displayed strong variations in their prevalence, both with respect to each other, as well as with respect to the age-group and Cohort-Type of the study-cohorts. Many of these variations replicated previous observations. The pattern of association of the non-Bifidobacterial community with the representation of the different Bifidobacterial species strongly correlated with their prevalence rates across study-cohorts of different age-groups and Cohort-Type. (Right Panel) Heatmap showing, for each cohort, whether significant associations (P ≤ 0.05; purple) were detected between non-Bifidobacterial community composition and nine Bifidobacterial properties using PERMANOVA utilizing either Bray–Curtis or Kendall distances as the non-Bifidobacterial community beta-diversity measures.

Notably, we observed significant variations in the Bifidobacterial prevalence patterns depending upon the profiling strategies (16S-v/s-WGS) of the study-cohorts (PERMANOVA R-squared=0.14, P=0.001, **Figure S1F)**, with multiple taxa like *B. longum/adolescentis/catenulatum/animalis* showing significantly lower detection in 16S cohorts as compared to WGS (**Figure S1D**; **Figure 1**). These observations could be a reflection of previously reported methodological limitations of the 16S method that impede efficient profiling of Bifidobacterial amplicons^20^.

Next, we asked whether the variations in the resident gut microbiomes (specifically the non-Bifidobacterial taxa constituting the gut microbiomes) had a significant association with the overall detection of Bifidobacterial species as well as with the representation of the eight major Bifidobacterial taxa. For this purpose, in each cohort, we investigated whether the variations within the non-Bifidobacterial community across the corresponding gut microbiomes (using two different measures, Bray-Curtis and Kendall) were significantly associated with the representation of the Bifidobacterial species (and the total detected Bifidobacteria) (See **Methods**). Variations in the non-Bifidobacterial gut microbiome were significantly associated with the Bifidobacterial detection rates in 74% and 64% of the study-cohorts, when the community variations were profiled using the Kendall and Bray-Curtis measures, respectively (**Figure 1**). Consistency of associations varied across species and age-groups, with the consistency of associations in general decreasing from infant > adults > senior (**Figure S2A-B**). Five major Bifidobacteria (*B. longum/breve/bifidum/dentium/breve*) showed significant associations with non-Bifidobacterial gut community in >50% of the infant cohorts, and four of these (except *B. dentium*) showing the similar patterns also in the adult cohorts and decreasing in the senior cohorts. These patterns putatively indicate that the strength of the associations of the Bifidobacteria with the non-Bifidobacterial decreases with age (despite the prevalence patterns of the different species being relatively similar between the adult and senior cohorts; **Figure S1C**). Among the species, *B. longum*, (showing the highest prevalence across study-cohorts) also exhibited the most consistent associations with the non-Bifidobacterial community across all age groups. This was followed by *B. bifidum*, *B. pseudocatenulatum*, and *B. breve*, with *B. breve* and *B. bifidum* much stronger associations in the infant cohort.

These patterns suggest that certain non-Bifidobacterial lineages may either influence, or be influenced by, the presence of specific Bifidobacterial taxa. To investigate this further, we analyzed association-patterns between non-Bifidobacterial taxa and both the prevalence and abundance of eight major *Bifidobacterium* species across diverse cohorts.

### Non-bifidobacterial taxa exhibit distinct association-patterns with Bifidobacteria, varying by subject age, lifestyle (industrialization), and microbiome profiling method

Our objective here was to identify which non-Bifidobacterial taxa showed consistent association with the individual with the overall *Bifidobacterium* prevalence and the abundances of the eight major Bifidobacterial species across gut microbiomes and how did these association-patterns vary in subjects from different age-groups (Infant/Adult/Senior) and life-style patterns (IndustrializedUrban/UrbanRuralMixed/RuralTribal). We performed these in three steps.

First, we removed sparsely detected taxa across gut microbiomes (described in **Text S1**; **Figure S3**). Removal of such taxonomic ‘noise’ is necessary as they can drastically decrease the sensitivity of the subsequent association analyses. Next, for each Bifidobacterial feature, we identified the sets of non-Bifidobacterial species-level-taxa, consistently identified as the ‘top-predictors’ across the study-cohorts. For this, we built study-cohort-specific Random Forest (RF) models using subsets of consistent gut-associated taxa (identified in the first step) to predict the corresponding Bifidobacterial features and selected consistent top-predictors across each model (See **Methods**; **Tables S2–S4**).

Different non-Bifidobacterial taxa showed varying and cohort-specific predictive power for different Bifidobacterial features, as indicated by their feature importance scores (**Tables S2–S5**; **Figure 2**; **Figures S4–S11**). These patterns varied by profiling method (16S/WGS), age-group (Infant/Adult/Senior), and lifestyle (Industrialized Urban/Others). For example, for overall *Bifidobacterium* detection across age groups, while some non-Bifidobacterial taxa were consistently identified as ‘top-predictors’ across all cohorts of an age-group, others being specific to either 16S or WGS datasets (**Figure 2**). There were also clear differences top-predictor taxa lists and their feature importance scores across infant, adult and senior cohorts. RuralTribal cohorts displayed distinct feature importance patterns compared to IndustrializedUrban cohorts, with mixed cohorts showing intermediate profiles. Feature importance patterns of non-Bifidobacterial taxa for predicting the abundance of the eight Bifidobacterial species also showed strong cohort-specific associations (**Figures S4-S11**). These patterns were significantly linked to sequencing type, age group, and lifestyle across 7, all 8, and 4 Bifidobacterial-species, respectively (**Figures S12–S14**). RF models revealed that specific non-Bifidobacterial taxa were strongly predictive of various Bifidobacterial traits. However, the strength of these associations differed drastically with cohort characteristics— sequencing method, age group, and lifestyle—with cohorts sharing similar traits exhibiting more similar association-patterns.

**Figure 2.**
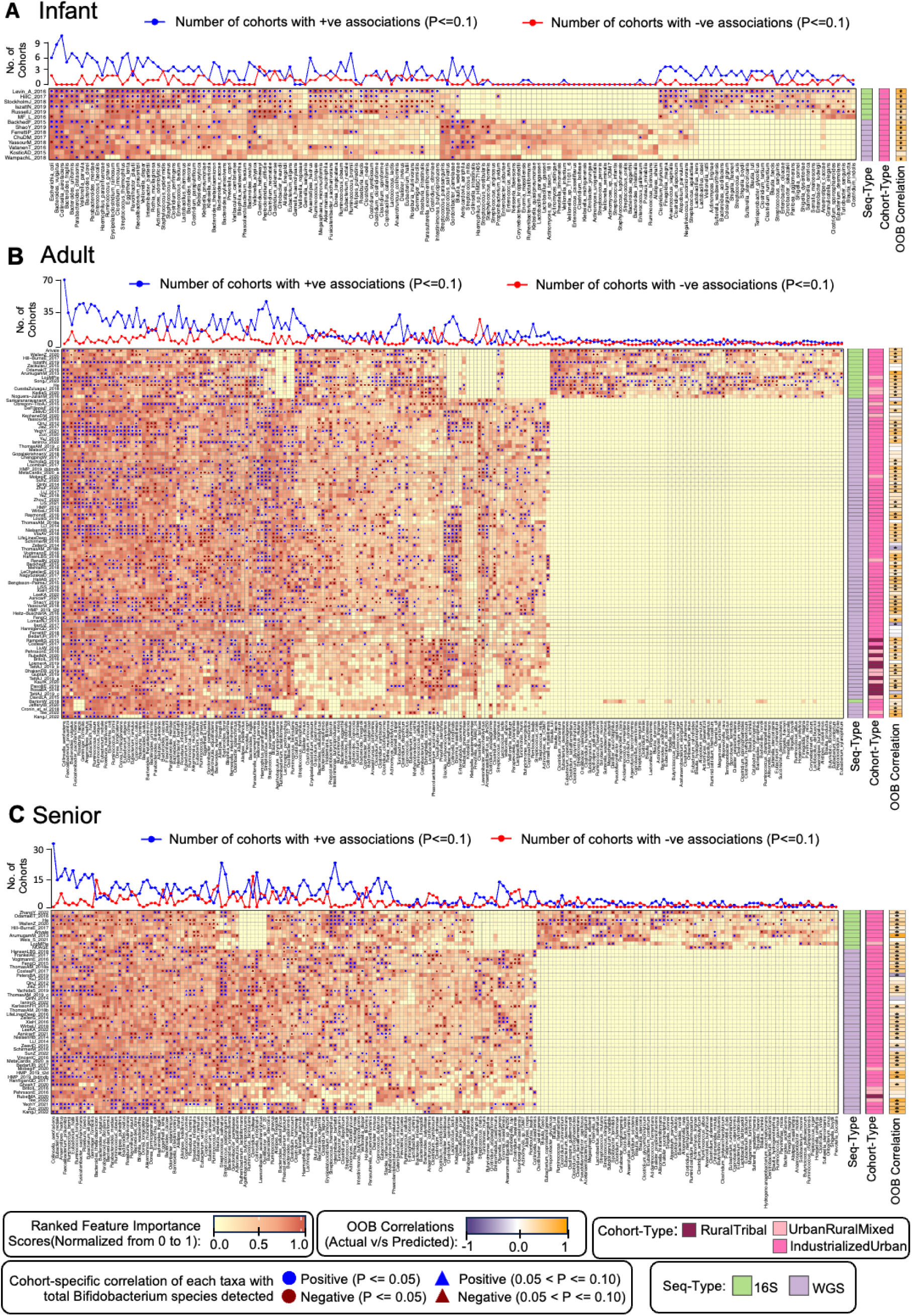
Random Forest (RF) models identified distinct sets of non-Bifidobacterial taxa that consistently associated with overall *Bifidobacterium* detection patterns, serving as top predictors of the total number of Bifidobacterial taxa detected in the gut microbiome. These associations varied across cohorts stratified by profiling method (16S/WGS), age group (Infant/Adult/Senior), and lifestyle (especially between IndustrializedUrban and RuralTribal groups). A-C. Heatmap showing the variation of the rank-normalized feature importance scores of the top predictor taxa for the Random Forest models built for Infant (A), Adult (B) and Senior (C) cohorts. The profiling strategy (Seq-Type), and the life-style (Cohort-Type) of each cohort is indicated as strips as shown for each heatmap. Also indicated are the out-of-bag (OOB) correlations between the actual and RF-predicted total number of *Bifidobacterium* species for each cohort. Symbols within the heatmap cells indicate both the direction (positive or negative) and strength (based on p-values) of the association between each non-Bifidobacterial taxon (columns) and Bifidobacterial detection within each cohort (rows). Line plots above each heatmap summarize these associations and show the number of cohorts where each non-Bifidobacterial taxon displays significant positive or negative associations with total Bifidobacterial species (p ≤ 0.1). The extent and consistency of these associations varied by the specific non-Bifidobacterial taxon, with some showing notably higher predictive power and consistent directionality. Association patterns were also cohort-dependent, namely Infant cohorts exhibited distinct sets of associated taxa compared to other age groups (same for other age-groups); 16S and WGS cohorts showed differing patterns; and RuralTribal cohorts displayed consistent association patterns that were distinct from those observed in IndustrializedUrban cohorts. These patterns were captured in the Association-Scores as described in Figure 3. We conducted similar analyses to identify non-Bifidobacterial taxa consistently associated with the abundances of the eight major Bifidobacterial species (results summarized in **Figures S4–S11**).

Finally, we investigated the directionalities of these associations. For a given Bifidobacterial trait and cohort-sets of specific characteristics (sequencing-type: 16S/WGS; age-group: Infant/Adult/Senior and cohort-lifestyle: IndustrializedUrban/UrbanRuralMixed/Rural), we selected the corresponding set of top-informative (top-predictor) non-Bifidobacterial taxa (listed in **Tables S2-S4**; **Figures S4-S11**), and then computed the Spearman correlations between the abundances of these taxa and the corresponding Bifidobacterial trait across all study-cohorts of that type (**Tables S5-S10**). These Spearman correlation patterns were then utilized to compute Association-Scores of the different non-Bifidobacterial taxa with different Bifidobacterial traits for the given cohort-type (**Figure 3A**). The Association-Scores were quantitative measures of the extent of association of each non-Bifidobacterial taxa with each Bifidobacterial feature in microbiome cohorts with specific characteristics, stratified based on cohort-characteristics (profiling strategy, age-group and cohort-lifestyle).

**Figure 3.**
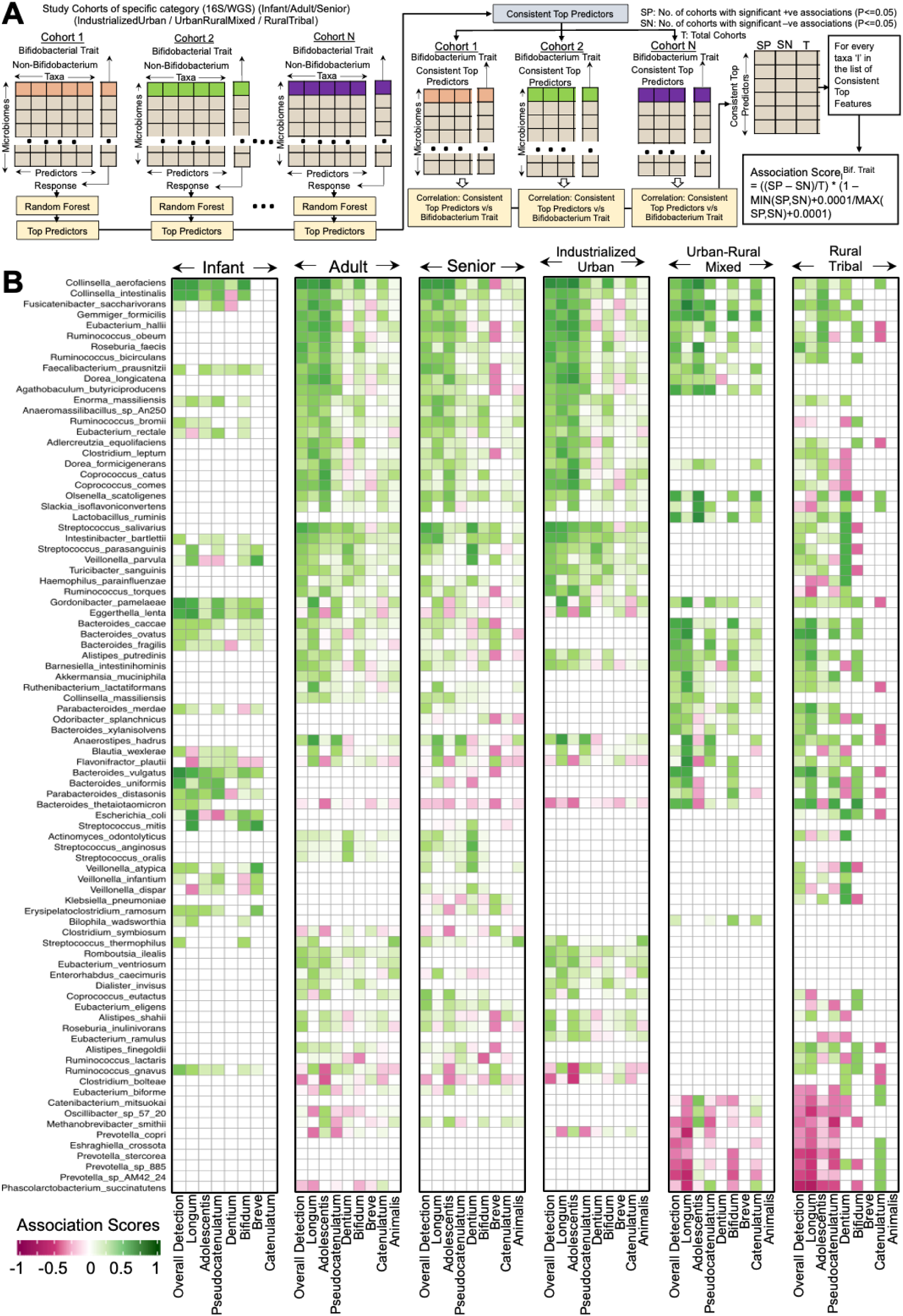
Association patterns of the non-Bifidobacterial taxa with the different Bifidobacterial properties could be quantitatively captured as Association-Scores, stratified by profiling-strategy or Seq-Type (16S/WGS), age-group (Infant/Adult/Senior), and lifestyle category (IndustrializedUrban/UrbanRuralMixed/RuralTribal). **A.** Schematic illustration of the methodology used to compute the Association-Score for a given non-Bifidobacterial taxon with a specific Bifidobacterial trait within cohorts of a defined category. **B.** Association-Score patterns for different non-Bifidobacterial taxa with various Bifidobacterial traits across cohorts of different categories. Only those non-Bifidobacterial taxa with showing consistent association patterns (Association-Score >= 0.25) with the different Bifidobacterial traits in at least two *Bifidobacterium*-trait-Cohort-type combinations are shown here. The complete Association-Scores list and the Association-Scores for 16S cohorts are provided in **Tables S11-S12**.

Analysis of these Association-Scores revealed several notable patterns. Two *Collinsella* species (*C. aerofaciens/intestinalis*) and *Faecalibacterium prausnitzii* consistently showed high Association-Scores with multiple Bifidobacterial taxa across all age groups and lifestyles (**Figure 3B**). Association-patterns for other non-Bifidobacterial taxa differed by age and lifestyle. In infant gut microbiomes, taxa such as *Eggerthella lenta* and *Gordonibacter pamelaeae* (both Actinobacteria phylum, like *Bifidobacterium*), along with *Bacteroides vulgatus/uniformis*, *Parabacteroides distasonis*, and *Erysipelatoclostridium ramosum*, exhibited high Association-Scores with several Bifidobacteria. However, these associations weakened or disappeared in adult and senior microbiomes.

In contrast, adult and older person gut microbiomes showed consistently high Association-Scores between multiple Bifidobacterial taxa and several butyrate-producing Firmicutes genera, including *Coprococcus*, *Eubacterium*, *Ruminococcus*, *Roseburia*, *Gemmiger*, *Dorea*, and *Fusicatenibacter,* that are linked to health benefits in previous studies^16,21,22^. Additionally, high Association-Scores were observed for oral-associated taxa such as *Streptococcus*, *Veillonella*, and *Haemophilus*. Notably, these scores were stronger in adults but declined in seniors.

Association-Scores also differed along the lifestyle/industrialization-gradient from IndustrializedUrban to RuralTribal gut microbiomes. In RuralTribal cohorts, Association-Scores with Firmicutes were generally reduced, except for *Fusicatenibacter saccharivorans*, but were consistently high for several Bacteroidetes taxa, e.g. *Bacteroides caccae/ovatus*, *Alistipes putredinis*, *Odoribacter splanchnicus*, *Barnesiella intestinihominis*, and uniquely, *Akkermansia muciniphila*. A consistent feature of RuralTribal cohorts was negative Association-Scores for *Prevotella* species (*copri, stercorea, sp. AM.42_24, sp. 885*), *Catenibacterium mitsuokai*, *Phascolarctobacterium succinatutens*, *Oscillibacter sp. 57_20*, and *Eubacterium biforme*. UrbanRuralMixed cohorts showed intermediate patterns, with positive associations for butyrate-producing Firmicutes (as seen in IndustrializedUrban cohorts) alongside elevated scores for *Alistipes*, *Bacteroides caccae/ovatus*, *Odoribacter*, and *Barnesiella*, and consistent negative associations for *Prevotella*, *Catenibacterium*, *Oscillibacter*, and *Phascolarctobacterium*, resembling patterns observed in RuralTribal groups. We computed the Association-Scores separately for the 16S cohorts (**Tables S11-S12**). Many of these patterns were also reproduced across the Association-Scores for each age-group and cohort-lifestyle sub-groups were reproduced across 16S and WGS gut microbiomes. Despite having variations in the detection of specific taxa, we observed significant correlations between the Association-Scores computed for the commonly occurring taxa between the 16S and the WGS datasets (**Figure S15**). Across the 32 combinations involving Bifidobacterial species (and overall Bifidobacterial detection) and the four different cohort categories (infant, adult/senior, adult-IndustrializedUrban, adult-OtherNonIndustrialized), for 25 combinations (∼78%) we observed a significant positive correlation between the Association-Scores for the different non-Bifidobacterial taxa obtained for WGS and 16S study-cohorts. We next investigated the variability of these association-patterns across different disease groups.

### Association-patterns are subtly altered in different diseases with specific disease groups showing similar non-Bifidobacterial associations with different *Bifidobacteria*

Given the limited availability of patient gut microbiomes of different diseases from non-Industrialized populations, here, we specifically investigated the variation of non-Bifidobacterial association-patterns (with different Bifidobacterial properties) in the adult/senior gut microbiomes from IndustrializedUrban cohorts. To investigate this in a comprehensive manner, we further supplemented our initial global gut microbiome collection (of 45,809 gut microbiomes, 121 study-cohorts) with data from 27 study-cohorts (an additional 5,249 gut microbiomes) containing matched diseased and non-diseased gut microbiomes (resulting in a total of 15,095 gut microbiomes from 69 study-cohorts with matched-diseased-control sub-groups) (See **Methods**, **Text S2** and **Table S1**). For ensuring robustness of findings, we focused on nine diseases, namely, Colorectal Cancer (CRC), Polyps, Prediabetes, Type-II-Diabetes (T2D), Cardiovascular Disease (CVD), Inflammatory Bowel Disease (IBD), COVID, Parkinsons and Liver-Disease, for which > 3 study-cohorts were available in our data collection. For each of these diseases, we computed the Association-Scores of the non-Bifidobacterial taxa with the different Bifidobacterial traits as described in the previous section (**Figure 3A**; **Methods**), and compared the same with those obtained with respect to the non-diseased controls from the same study-cohorts (**Figure 4**).

**Figure 4.**
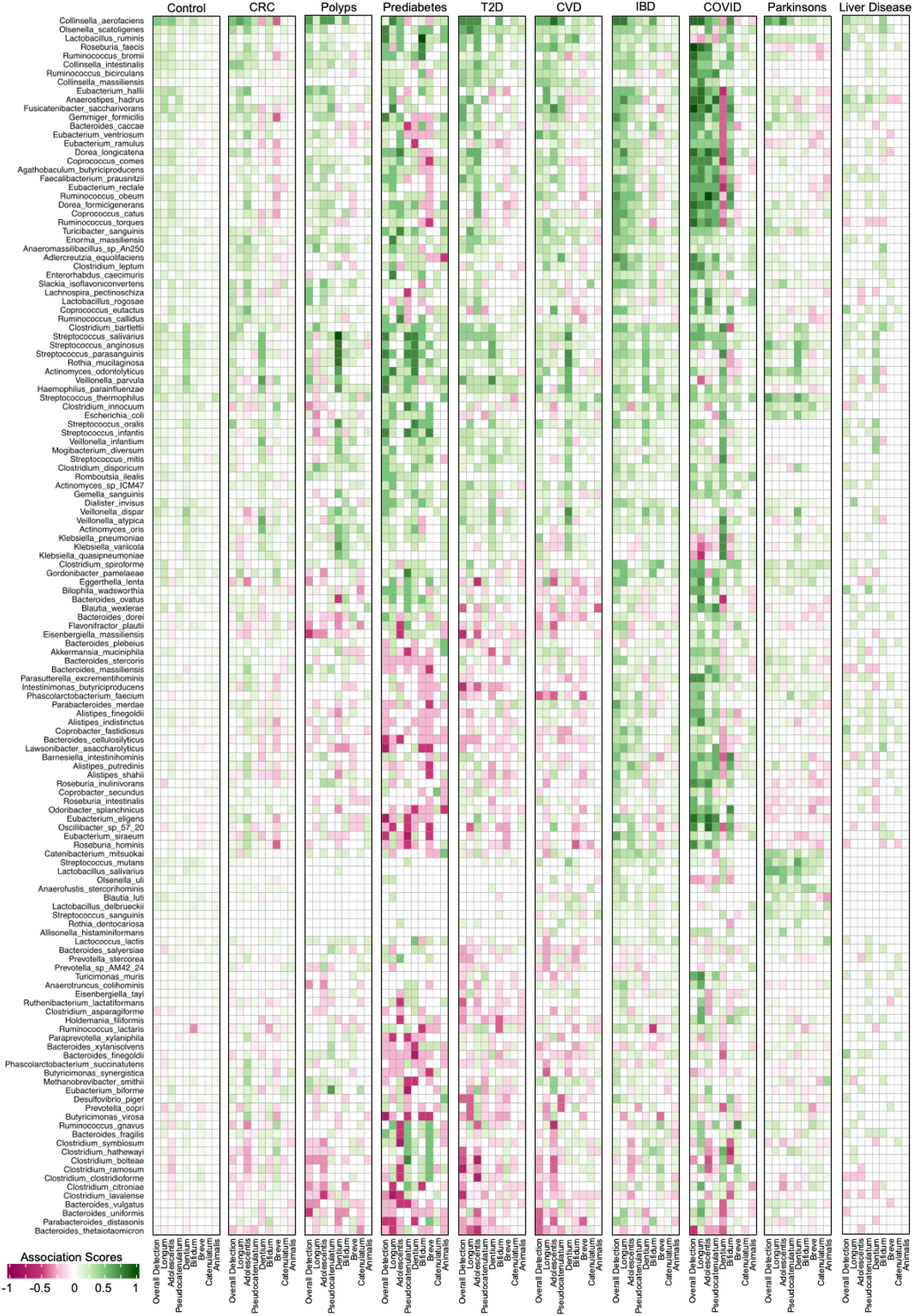
Association patterns of the non-Bifidobacterial taxa with the different Bifidobacterial properties show specific variations not only between diseased and non-diseased controls, but also across the nine different diseases. Association-Scores computed for different non-Bifidobacterial taxa with various Bifidobacterial traits across cohorts of nine different diseases viz. Colorectal Cancer (CRC), Polyps, Prediabetes, Type-II Diabetes (T2D), Cardiovascular Diseases (CVD), Inflammatory Bowel Disorder (IBD), COVID, Parkinson’s Disease, and Liver Disease) and apparently non-diseased individuals obtained by investigating the matched-control-sub-cohorts for each of the 69 studies corresponding to these nine diseases. Only those diseases were considered where the data was available for more than three study-cohorts and only those non-Bifidobacterial taxa showing consistent association patterns (Association-Score >= 0.25) with the different Bifidobacterial traits in more than two Bifidobacterium-trait-Disease combinations are shown here. The colour scale indicates the range of association score spanning from -1 to 1.

For most Bifidobacterial traits (except *B. catenulatum/animalis* abundance) and most diseases (excluding liver disease and Parkinsons’), the Association-Scores of non-Bifidobacterial taxa with Bifidobacterial traits in non-diseased controls showed significant positive correlations to those observed for the diseases indicating overall similar association-patterns (**Figure 4**; **Figure S16**). These similarities included consistent positive associations with *Collinsela*, multiple butyrate-producing members (as observed earlier; **Figure 3B**) (especially with overall detection and *B. longum/adolescentis/pseudocatenulatum* abundances). There were also specific variations between non-diseased and diseased gut microbiomes. Unlike the non-diseased sub-group, almost all diseases were characterized by consistent negative associations of multiple Bifidobacterial traits with multiple lineages like, pathobiont *Clostridium* (*C. bolteae/citroniae/ramosum/asparagiforme/hathewayi*), *Ruminococcus gnavus* and *Bacteroides fragilis*, that have been positively associated with frailty, inflammation and multiple diseases across studies^21,23^ (**Figure 4**). Consistent positive associations with butyrate producers and negative associations with disease-enriched pathobionts were similarly observed across 16S and WGS cohorts for four of these diseases (with enough study-cohorts available) (**Figure S17**). These further highlight the health-associated aspect of *Bifidobacteria* (**Figure 4**).

There were also notable variations amongst different diseases. Cardiometabolic diseases like prediabetes, T2D and CVD were characterized by consistent negative associations of multiple Bifidobacterial properties with specific *Bacteroides* (*B. stercoris/massiliensis/cellulosilyticus*), *Roseburia* and *Alistipes* species along with *Lawsonibacter asaccharolyticus*, *Barnesiella intestinihominis*, *Odoribacter splanchnicus*, etc. In contrast, for gut-inflammation/infection-associated diseases (IBD and COVID), the patterns were reversed with these lineages showing consistent positive associations with multiple Bifidobacterial traits (**Figure 4**).

### Post-treatment increase of *Bifidobacterium* probiotics can be significantly predicted by microbiome-level receptive-scores, computed using the Association-Scores and baseline abundances of the administered *Bifidobacterium* probiotic

We next investigated the reproducibility and the translational applicability of these Association-Scores. We collated data from eight independent trials in this sub-analysis, encompassing a total of 1063 gut microbiomes from 442 subjects (**Table S1; Table S13;** described in detail in **Text S3**)^11,15,24–29^. While seven of these trials directly administered different live *Bifidobacterium* probiotics (either alone or in combination), one of the trials had used galacto-oligosaccharide (GOS), a prebiotic aimed at increasing Bifidobacterial species composition in general in infants.

Majority (5) of these eight trials-cohorts involved adult/senior individuals from IndustrializedUrban populations, with microbiome being profiled using WGS approach^11,24–26,28^. To investigate the replicability of the association-patterns of the non-Bifidobacterial taxa with the different Bifidobacterial properties, we recomputed their Association-Scores in this subset of five intervention cohorts and compared these scores with those obtained for the cohorts of similar types (Adult/Senior-IndustrializedUrban-WGS) in discovery cohorts (See **Methods**; **Text S4**; **Figure 5A**; **Table S14**). For each of the eight Bifidobacterial taxa, the association-patterns of the non-Bifidobacterial taxa were reasonably reproduced (significant positive correlations between 0.22-0.67) (**Figure 5A)**. Additionally, one of the cohorts involved infants from an IndustrializedUrban cohort (microbiome profiled using 16S)^15^. Here, we computed the correlations of the abundances of the different non-Bifidobacterial taxa with that for each of the seven Bifidobacterial taxa (except *B.animalis*). Subsequent correlation of these associations with the corresponding discovery-cohort-obtained Association-Scores in study-cohorts of this category (Infant-IndustrializedUrban-16S), also showed significant positive correlations (ranging from 0.27 to 0.55) (**Figure 5A**; **Table S15**). Thus, cohort-type-stratified association-patterns of the non-Bifidobacterial taxa with each of the Bifidobacterial species are robust and significantly reproducible.

**Figure 5.**
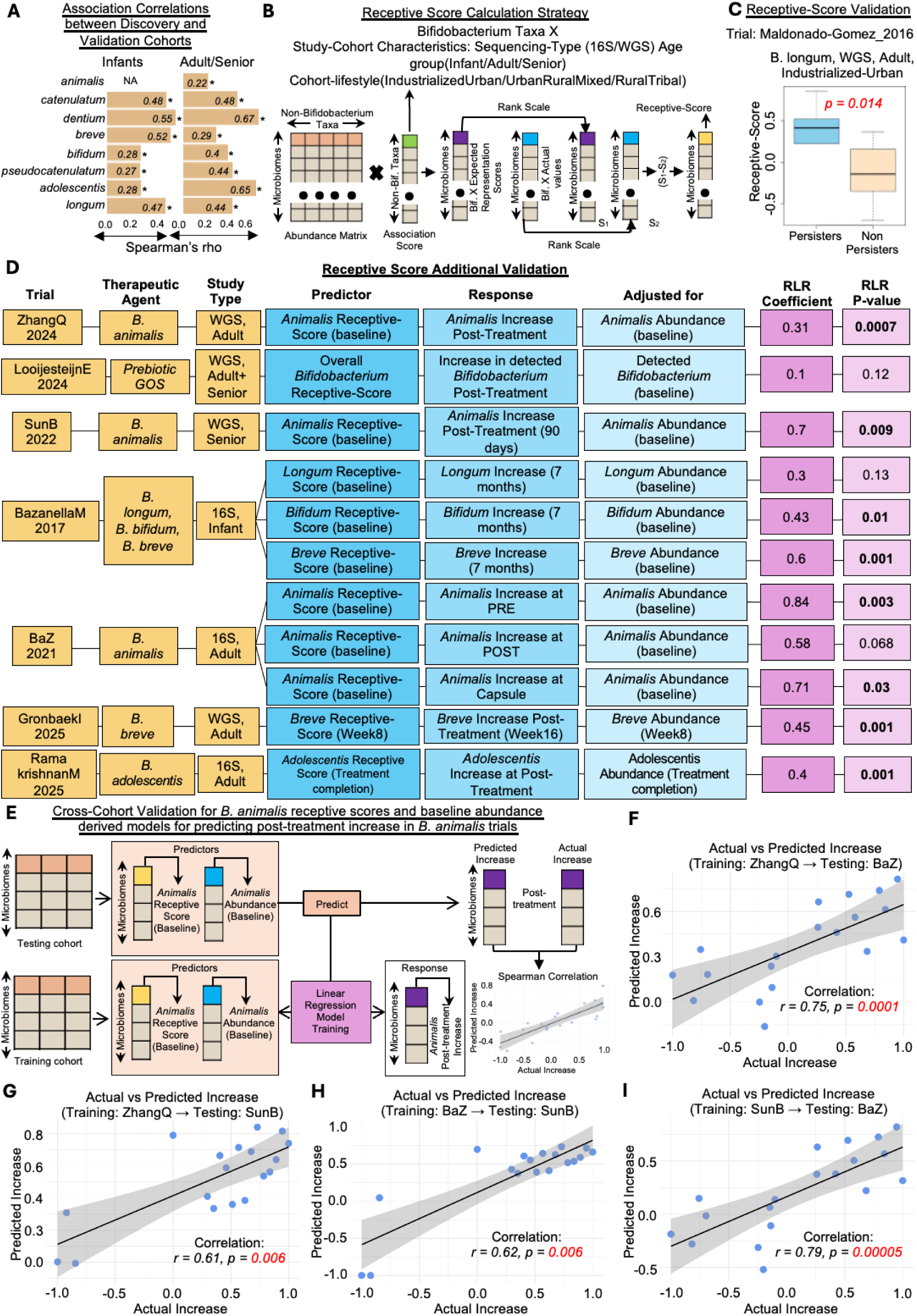
Reproducibility of Association-Scores and the utility of baseline *Bifidobacterium* Receptive-Scores for personalized prediction of future increase/persistence. A. (left) Bar-plots showing the Spearman-Correlations between the Association-Scores of different non-Bifidobacterial taxa with the abundances of seven Bifidobacteria (in the infant-IndustrializedUrban-16S cohorts of the discovery datasets) and their abundance correlations with the corresponding Bifidobacteria in the infant/new-born-associated Bazanella *et al* cohort validation dataset. (right) Bar-plots showing the Spearman-Correlations between the Association-Scores of the different Bifidobacterial taxa with the eight major Bifidobacteria computed for the WGS-profiled Adult/Senior cohorts with IndustrializedUrban population-type in the discovery dataset with the same Association-Scores re-computed within the cohorts of the same microbiome-type in the validation dataset. Overall, these results indicate a strong and significant reproducibility of the association patterns of the different non-*Bifidobacterium* taxa with the different *Bifidobacterium* species in the discovery and validation cohorts. **B.** Strategy for computing the microbiome-level Receptive-Score for a given *Bifidobacterium* property, given the microbiome-type of the specific cohort (characterized by the specific age-group, cohort-life-style and microbiome-profiling strategy). **C.** Boxplots showing the significantly higher baseline *B. longum* Receptive-Scores for the Persisters compared to the Non-Persisters in the Maldonado-Gomez *et al* cohort, along with the Mann-Whitney-test p-value. **D.** Horizontal flow diagrams illustrate significant associations between baseline Receptive-Scores and post-treatment increases in target *Bifidobacterium*-associated properties. Post-treatment increase is defined as the rank-scaled difference between immediate post-treatment and the baseline values. Associations were assessed using RLRs (Methods), with Receptive-Score as the predictor, post-treatment increase as the outcome, and baseline property levels as confounders (specified per model). Each diagram presents results for seven *Bifidobacterium*-focused trials, covering eleven ‘trial-Bifidobacterium-property’ scenarios. The ‘RLR Coefficient’ column signifies positive association and ‘RLR P-value’ indicates significance of the association. **E.** Approach adopted for the cross-cohort validation using the linear regression model designed to predict post-treatment increase of *B. animalis* based on Receptive-Scores and baseline abundance of *B. animalis*. **F-I:** Results of the cross-cohort validation of these linear regression models with SunB and BaZ as the testing cohorts.

We next investigated the translational applicability of our findings. In other words, we explored whether the cohort-type-stratified Association-Scores of the various non-Bifidobacterial taxa for the different *Bifidobacteria* (**Figure 3A**) could be utilized to formulate microbiome-level quantitative measures for stratifying or ranking individuals based on their gut microbiome’s receptivity to different *Bifidobacterium* ‘probiotics’. We referred to these measures as microbiome-level ‘Receptive-Scores’. Given a target *Bifidobacterium* taxon (or the overall *Bifidobacterium* detection) and microbiomes of a specific category (based on host-age, host-life-style and 16S/WGS profiling), the corresponding Receptive-Scores were computed using a two-step approach (**Figure 5B**). In the first step, given a set of microbiomes of a specific category, we considered the abundances of all non-Bifidobacterial taxa and computed cumulated weighted sums, where in the abundances of the non-Bifidobacterial taxa were weighted by their corresponding Association-Scores computed previously for microbiome-types of that category (**Figure 3B**) (the abundances of only those non-Bifidobacterial taxa with matching Association-Scores were considered). This approach ensured that microbiomes with a greater abundance of non-Bifidobacterial taxa which globally show consistent positive associations with a target *Bifidobacterium* received higher scores (and those having greater abundance of microbes showing consistent negative associations received lower scores). Ranking these scores across a population-cohort facilitates effective ranking of individuals based on how well their current microbiome is expected to support the target *Bifidobacterium* species. We referred to this the Expected-Representation-Score for that *Bifidobacterium*.

Expected-Representation-Scores quantify how supportive a community is for a given *Bifidobacterium*, based on the composition of other resident taxa. However, the receptivity of a community for a particular *Bifidobacterium* taxon (in terms of the future success of its administration/supplementation in a community, measured by persistence or increase) also depends on the existing abundance of that *Bifidobacterium*. Communities already rich in a specific Bifidobacterium may have limited capacity to accommodate more members of the same lineage and might not show a noticeable increase/persistence after its administration^30,31^. Therefore, it’s equally important to rank individuals based on the current (or the baseline) abundance of the target *Bifidobacterium* in their gut. Thus, for every gut microbiome from a specific population group (based on cohort-characteristics as described above) and a target *Bifidobacterium* taxon, we can calculate a Receptive-Score as the difference between the ranked Expected-Representation Score and the ranked baseline abundance of the target *Bifidobacterium* taxon. This score indicates how well an individual’s gut microbiome is receptive to support an increase in the target *Bifidobacterium* after administration. A higher Receptive-Score suggests that the microbiome can accommodate a higher representation of the target *Bifidobacterium* probiotic. It identifies individuals whose gut environment is conducive to increased abundance of the target *Bifidobacterium*, considering both its current levels and the supportive microbial context.

Next, we validated the concept/applicability of Receptive-Scores in these eight validation *Bifidobacterium* intervention trials (**Table S13**). The first trial to be investigated was by Maldonado-Gomez *et al*, which had investigated the persistence of *Bifidobacterium longum* AH1206 in 23 individuals^11^. The authors had observed that, in only 30% individuals (the ‘Persisters’), the strain showed long-term persistence and that this pattern was associated with variations in the baseline microbiome. We thus checked whether the Receptive-Scores for *B. longum* (using the corresponding *B. longum* Association-Scores in microbiome types profiled using WGS in adult individuals with IndustrializedUrban life-styles) in the baseline gut microbiomes of this specific trial dataset, could distinguish future ‘Persisters’ and ‘Non-Persisters’. Notably, the computed baseline Receptive-Scores were significantly higher for Persisters compared to Non-Persisters (Mann-Whitney U-test P-value: 0.014; **Figure 5C**). This provided the first validation, suggesting that baseline Receptive-Scores can effectively identify individuals likely to exhibit future persistence or an increase of the target probiotic.

Encouraged by this initial validation, we investigated seven additional *Bifidobacterium*-associated trials (See **Text S3**; **Table S13**)^15,24–29^. For each of the trials, we identified the target *Bifidobacterium*-related property (or properties) that was (were) expected to be increased. The property investigated was either ranked abundance of any of the eight *Bifidobacterium* taxa or the overall *Bifidobacterium* detection. To evaluate whether the baseline Receptive-Score for a given target property was predictive of its future change, we employed robust linear regression (RLR) models across each trial (see **Methods**; **Figure 5D**), where we tested we tested whether the Receptive-Score was associated with the direction and significance of the property’s post-treatment increase, after including the ranked baseline abundance as a covariate to control for its potential confounding effects. This was because both the Receptive-Score and the ranked post-treatment increase are derived from the ranked baseline abundance (**Figure 5D**). Across the 11 trial–Bifidobacterium–property combinations (in seven trials), the RLR-models showed significant positive relationship between the baseline Receptive-Score and an increase of the target Bifidobacterium(s) (RLR coefficient > 0; P-value <= 0.05) in 8 combinations. Thus, overall, 9 out of 12 trial-Bifidobacterium-property combinations (75%) (including Maldonado-Gomez^11^), we observed that higher baseline Receptive-Scores are significantly associated with increased post-treatment increase/persistence of the target *Bifidobacterium*.

In addition, three trials featured *B. animalis* as the administered probiotic, all conducted in IndustrializedUrban Chinese populations—two using whole-genome shotgun (WGS) sequencing and one using 16S rRNA gene sequencing^24,26,27^. We further assessed the robustness of the Receptive-Score framework using cross-cohort validation. Here, for each trial, we trained a cohort-specific linear-regression model to predict the increase in *B. animalis* at the immediate post-treatment time-point, using baseline Receptive-Score and baseline *B. animalis* abundance as predictors. Each trained model was then tested on the remaining two cohorts to evaluate its ability to predict *B. animalis* increase at the corresponding post-treatment time-point, based on the baseline abundance and Receptive-Scores in those cohorts (**Figure 5E**). Across each of the six training-testing cohort pairs, there was significant positive correlation between the actual and predicted increase in *B. animalis* in the testing cohorts (**Figure 5F-I**; **Figure S18**), indicating the translational applicability of the concept of Receptive-Scores to predict response in individuals across novel ‘unseen’ cohorts.

### Specific genome-level functional features of the different non-Bifidobacterial taxa link to their Association-Scores with the different Bifidobacterial lineages

Associations between microbes are reflections of their functional interactions, (including the phenomenon of cross-feeding) which may promote better colonization of a Bifidobacterium in a community^30,31^.

Using a systematic approach, we collected a set of 28,991 genomes belonging to 72 consistently occurring non-Bifidobacterial gut microbial taxa for which Association-Scores were obtained against multiple Bifidobacterial traits (**See Methods**; **Text S5; Figure 3B**; **Figure 6A; Figure S19**). We annotated these genomes using the eggNOG mapper pipeline (See **Methods**)^32^, to obtain genome-level functional signatures based on nine different functional schemas (See **Methods**; **Figure 6A; Figure S19**). We then grouped co-linear/redundant functional feature annotations (i.e. the same gene/protein annotated by multiple different annotations under multiple different schema) to finally obtain a list 16,077 non-redundant ‘functional-groups’ (**See Methods**).

**Figure 6.**
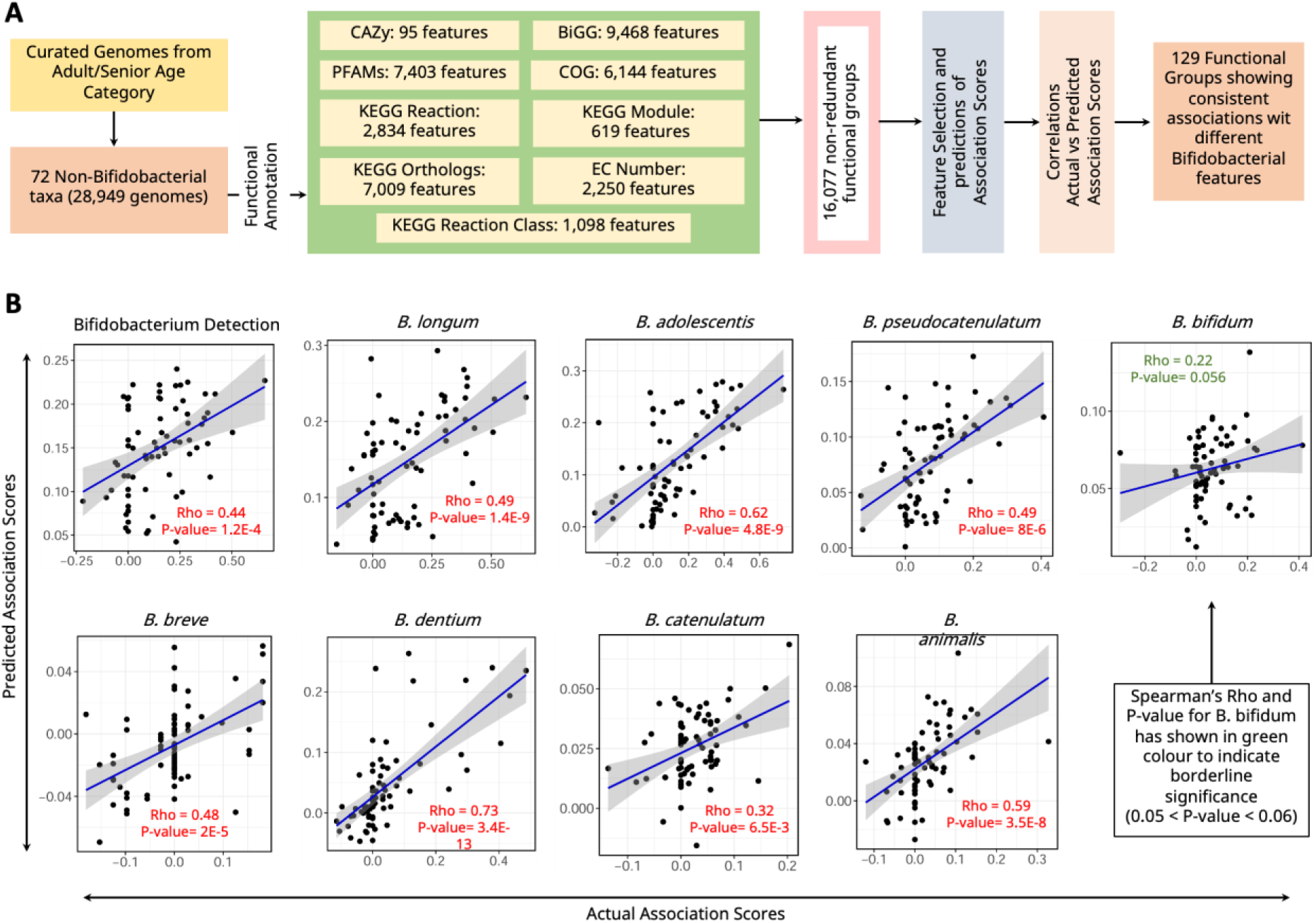
Associating non-Bifidobacterial taxa derived functions which show predictive ability of different Bifidobacterial Association-Scores. **A.** Schematic overview of generating non-Bifidobacterial taxa derived 129 functional groups showing consistent associations with Bifidobacterial taxa, starting from curation of high-quality genomes (including MAGs) followed by functional annotations, random-forest based feature selection, prediction of association scores and finally correlating original vs predicted association scores to identify the consistent set of functional groups **B.** Here in this panel the scatter plots show the actual vs predicted Association-Scores along with the best fit curve which shows the extent of positive correlation in our case. The Spearman’s rho and p-value have been shown in red colour (where p-value < 0.05) on the individual plots except for B. *bifidum* (green colour) which shows marginal significance with 0.05 < p-value < 0.06.

Subsequently, for each of the nine Bifidobacterial properties, we devised a two-step Random Forest (RF) machine-learning based approaches to systematically identify the set of more informative functional-groups for the prediction of the corresponding Association-Scores (See **Figure 6A; Figure S19; Methods**). These two-step RF models, for each *Bifidobacterium* property resulted in distinct sets of informative functional-groups and demonstrated strong predictive power for the prediction of the corresponding Association-Scores with significantly positive out-of-box correlations between the actual and predicted Association-Scores (ranging 0.32 to 0.73) for 8 of 9 properties (except *B. bifidum* Association-Scores which showed marginal significance: R: 0.22, P: 0.056) (**Figure 6B**). These results indicated that the Association-Scores of each non-Bifidobacterial taxa with respect to the different Bifidobacterial properties are strongly linked to their genome-level functions.

Applying further filtration criteria, we finally identified a set of 129 functional-groups that showed significant association with multiple *Bifidobacterium* properties (and were contributed by multiple non-Bifidobacterial taxa). The 129 functional-groups belonged to different functions ranging from ion-transport, adhesins, pili, carbohydrate transport/metabolism to energy metabolism and DNA replication and transcription (**Figure S20A**). Transmembrane proteins, transporters/exporters and cell-wall components along with cell communications and binding proteins constituted 35-38% of these functional categories (**Figure S20B-C**). We have provided a list of the non-Bifidobacterial protein sequences belonging to these functional groups in the github repository containing the results of this study (https://github.com/Sourav-MiRe/Project_Bifidobacterium).This list represents a set of non-Bifidobacterial functional genes that can be evaluated further to probe the intestinal ecology of commensal microbes.

## Discussion

The gut microbiome modulates responses to Mediterranean Diet interventions, medications, anti-cancer therapies and putative probiotics^33–36^. Microbiome-based therapies (including probiotics) have two different levels of responses, first, colonization/persistence/abundance-increase of the administered health-associated microbe(s) and second, the clinical efficacy. These may occur synergistically or independently. With regard to colonization/persistence/abundance-increase, a recent analysis of probiotic trials focused on cardiometabolic disorders, found that only ∼40% of trials showed a change in the gut microbiome composition^10^. Although variation of baseline gut microbiome has been implicated as a key factor in determining personalized colonization/persistence patterns of different ‘probiotic’ species^11,37^, systematic identification of the determinants (both taxonomic and functional) of colonization/persistence/high-representation patterns of health-associated taxa has been very limited.

Our study is an effort in this direction. This systematic investigation of > 50,000 gut microbiomes (148 cohorts), spanning diverse ages, lifestyles (industrialization-status-based), diseases and profiling strategies (16S, WGS), not only enabled the identification of different non-Bifidobacterial taxa showing consistent (positive or negative) association-patterns with the eight major Bifidobacteria, but also enabled assessment of their variability across different population-types. For example, a previous cross-cohort analysis on a smaller collection of 9,515 gut microbiomes identified *Collinsella aerofaciens* along with multiple butyrate-producing members of the Firmicutes clade to be associated with higher representation of *B. longum*/*adolescentis*. Here, we show that the positive associations with other butyrate producers is strong only in the adult gut microbiomes of the IndustrializedUrban cohorts. These association-patterns become less consistent in seniors and UrbanRuralMixed cohorts and absent in infants and RuralTribal cohorts. Equally notable were the variations in these association-patterns across different diseases. Diseases in general not only showed subtle variations in the way non-Bifidobacterial community associate with the different Bifidobacteria (with respect to a non-diseased state), but also cardiometabolic diseases (like Prediabetes, T2D, CVD) show distinct variations in these association-patterns as compared to those linked to inflammation/infection (IBD and COVID). By investigating a large collection of diverse gut microbiomes, our analysis extends the understanding of baseline-gut-microbiome-to-Bifidobacterial association-patterns.

A highlight of this study is the formulation of microbiome-level Receptive-Scores for different *Bifidobacterium* taxa, which emphasizes the translational applicability of our findings. Using a combination of cohort-type-specific patterns (i.e. Association-Scores) of the different non-Bifidobacterial taxa, their abundance in a given gut microbiome and the current abundance of the administered *Bifidobacterium* taxa, we formulated Receptive-Scores to quantify how well a given microbiome can facilitate persistence and accommodate more members of the administered Bifidobacteria. As proof-of-concept, we validated the concept of Receptive-Score in 8 independent trials, where baseline scores predicted post-treatment Bifidobacterium increase/persistence in 75% of cases. Using cross-cohort regression, we further show that combining baseline Receptive-Scores with Bifidobacterium abundance enables accurate prediction of colonization response, supporting patient stratification for probiotic interventions.

We additionally identify specific genome-level functional signatures of the different non-Bifidobacterial taxa that strongly correlated with their Association-Scores with the different Bifidobacteria taxa. These functional signatures (especially the 129 functional-groups showing consistent associations with the representation of multiple Bifidobacteria) indicate the putative mechanistic links of how different non-Bifidobacterial lineages can modulate the ability of the microbiome to support different Bifidobacterial lineages. Experimental investigation of these functional-groups can clarify the mechanistic bases of these interactions.

Given that it is data-driven investigation of publicly available microbiome data, the analysis has certain limitations. Age-group, life-style and profiling strategy are expected to predominantly influence the gut microbiome composition. However, the association-patterns between different Bifidobacteria and the non-Bifidobacterial lineages can be modulated by other host-associated factors like gender, diet and medications. We could not perform the stratification of association-patterns based on these metadata because of their unavailability in many of the included datasets. Secondly, despite inclusion of multiple study-cohorts across a life-style gradient and investigating association-patterns individually across different cohort-types, there were a dearth of study-cohorts from non-Industrialized populations especially in the infant and senior age-groups. Lastly, while we have investigated the association-patterns at the species-taxonomic-level, multiple Bifidobacteria have shown consistent strain-wise variations in their colonization patterns and thus may show variabilities of association-patterns with different non-Bifidobacterial lineages^38^. This forms the objective of one of our future investigations.

Despite these limitations, the identification of consistent association-patterns between non-Bifidobacterial and Bifidobacterial lineages stratified based on the cohort-type and microbiome profiling approaches, the utilization of these association-patterns to build microbiome-level Bifidobacterial Receptive-Scores and the robust validation of these scores to predict personalized patterns of persistence/increase of the administered *Bifidobacterium* taxa in eight diverse intervention trials, demonstrate the applicability of our global analysis in formulation of approaches to predict personalized responses to Bifidobacterial-probiotic interventions.

## Resource availability

### Lead contact

Further information and requests for resources and data should be directed to and will be fulfilled by the lead contact, Tarini Shankar Ghosh.

## Material availability

This study did not generate new unique reagents.

## Data and code availability

- This paper analyzes existing, publicly available data, accessible at the European Nucleotide Archives (ENA) and curated Metagenomic Data (see Methods). Study details are listed in Table S1.
- All original code has been deposited at https://github.com/Sourav-MiRe/Project_Bifidobacterium and is publicly available.
- Any additional information required to re-analyze the data reported in this paper is available from the lead contact upon request.

## Acknowledgments

T.S.G. acknowledges the Department of Biotechnology, Ministry of Science & Technology, Government of India for the Ramalingaswami Re-entry Fellowship (BT/HRD/35/02/2006). T.S.G acknowledges IIIT-Delhi for the Discovery Track Investigator grant. S.G. acknowledges IIIT-Delhi for the Institute-Fellowship. A.A. acknowledge the Department of Science and Technology (DST-INSPIRE) for the INSPIRE fellowship. The P.W.O’T. laboratory is supported by Science Foundation Ireland (APC Microbiome Ireland: 12/RC/2273_P2), the EU’s Horizon-Europe research and innovation programme (grant-agreement nos. 101079777 with the MicrobAIome consortium and; 101083671 with the Microbiomes4Soy consortium).

## Author contributions

Conceptualization, T.S.G.; methodology, formal analysis, validation, S.G., T.S.G., A.A., C.S.; investigation, S.G., T.S.G., A.A., P.W.O’T.; writing—original draft, T.S.G., S.G., A.A.; writing—review and editing, T.S.G., P.W.O’T., F.S., V.A.; supervision, T.S.G.; funding acquisition, T.S.G.

## Methods

### Collation of discovery study-cohorts

The collation of the datasets was performed integrating two sources (listed in **Table S1**). First was the dataset collation of 124 study-cohorts, as described in a recently published study from our group^16^. This group of studies majorly contained gut microbiomes from adult and senior populations. This was supplemented by 32 additional study-cohorts from infants/adult/senior available either as part of the curatedMetagenomicData repository (https://github.com/waldronlab/curatedMetagenomicData), or downloaded as part of extensive literature searches and processed for inclusion in the analysis^39–44^.

The individual microbiomes were categorized into three age-based groups according to the age of the corresponding hosts: infants/newborns (≤2 years), adults (18–60 years), and seniors (>60 years). In certain studies, while individual-level age data were unavailable, the study-cohorts were reported to fall within specific age ranges, as indicated in the respective publications^45–51^. In such cases, all microbiomes from those cohorts were assigned to the appropriate age category (infant, adult, or senior) based on the available cohort-level information. Additionally, each microbiome was assigned to one of three lifestyle groups, namely IndustrializedUrban, UrbanRuralMixed, or RuralTribal, that are reflective of the general lifestyle of the study-cohort. This cohort-lifestyle classification was performed using the nationality and geographic location of the participants using methodologies described in previous studies from our group^16,17^. For each study-cohort, we also recorded the profiling strategy employed, namely 16S rRNA gene sequencing (16S) or whole genome shotgun (WGS) sequencing. Additional metadata such as gender, disease or study condition, birth mode, and others were available for some cohorts. However, these variables were not included in the current analysis due to their limited availability across cohorts or because they have already been investigated in previous studies^12–14^.

### Taxonomic profiling

For datasets obtained using the curatedMetagenomicData repository, the taxonomic profiles of all shotgun datasets (or cohorts) generated using metaphlan3 were already available^52^. For this reason, to maintain uniformity of profiling, we profiled the taxonomic composition of all WGS microbiome datasets of cohorts of not obtained from curatedMetagenomicData using metaphlan3. Microbiome datasets using 16S were all profiled using SPINGO^53^. The taxonomic profiled obtained for each study-cohort were subdivided into sub-parts, one containing the abundances of the non-Bifidobacterial taxa (which was renormalized after removing the abundances of the different Bifidobacterial species), and second containing the abundances of only the Bifidobacterial species.

### Filtering out sparsely detected species

We divided the study-cohorts into two bins based on their sequencing type. For each bin, we selected a subset of taxa detected consistently across multiple cohorts using the strategy summarized below (and described in detail in **Text S1**).

### Profiling of different Bifidobacterial across study-cohorts

To first profile the prevalence pattern, the abundance matrix corresponding to the different Bifidobacterial species were first converted to a detection matrix to investigate cohort-specific prevalence patterns for the different Bifidobacterial species. In the detection matrix, a given *Bifidobacterium* species with a relative abundance >= 0.0001 was marked as 1 (or detected) and 0 (not-detected) otherwise. The threshold of 0.0001 was determined based on previous studies using simulated metagenomes, especially using the Metaphlan approach, which have observed that, for abundance values >= 0.0001, there was a linear relationship between the actual abundance values, indicating reliable taxonomic assignments. While the detection profiles were utilized for investigating the association-patterns of different *Bifidobacterium* species with different cohort-specific characteristics like Sequencing-Type, Cohort-Lifestyle (or Cohort-Type) and host age-group, the abundance matrix was utilized for investigating the association of the non-Bifidobacterial community members with the overall *Bifidobacterium* representation as well as with the representation of the eight major *Bifidobacterium* species.

### Association of the detection pattern of different Bifidobacteria with the cohort Sequencing-Type, Cohort-Type and host age-group categories

To investigate the association-patterns of the eight Bifidobacterial species with different study-cohort-specific characteristics, we utilized linear regression models. We utilized three different regression models for this purpose, each with a specific objective and investigating the effect of a specific cohort characteristic after adjusting for the effects of all others. This is described in **Text S6**.

### Association of the overall non-Bifidobacterial beta-diversity with Bifidobacterium representation across study-cohorts

We utilized the PERMANOVA approach for this purpose. For a given *Bifidobacterium* species ‘i’ in a study-cohort ‘x’, we first computed the non-Bifidobacterial beta-diversity by computing the all-v/s-all sample distances using the renormalized relative abundance matrices of the non-Bifidobacterial taxa across the samples. This was performed using the Bray-Curtis and Kendall distance matrices and provided a quantitative measure of how different sample was respect to all the other samples in the cohort ‘x’ when considering only the non-Bifidobacterial taxa. Subsequently these variations were correlated with the abundance differences of Bifidobacterial species ‘i’ using PERMANOVA. This was performed using the adonis2 function of the vegan R package (v2.6-4). Significant association obtained using PERMANOVA indicates that variation of the species ‘i’ across the different samples is strongly correlated with community-composition of the non-Bifidobacterial taxa indicating a relationship between the two.

### Study-specific Random Forest models to identify the top predictive non-Bifidobacterial taxa for the different Bifidobacterial features across the cohorts

For each study-cohort, we built Random Forest based Machine-learning models for the prediction each of the Bifidobacterial features (overall *Bifidobacterium* detection and the abundances of the eight Bifidobacterial species), using the abundance profiles of the cohort-type-specific lists of non-Bifidobacterial taxa identified in **Text S1**. For each model (formulated for a given Bifidobacterial feature in a given cohort), we identified the top predictor non-Bifidobacterial taxa in the following manner. Taxa were sorted in decreasing based on their feature importance scores. Based on this ordering, for any set of top ‘I’ taxa, we computed the fractional cumulative importance (FCI) value as:

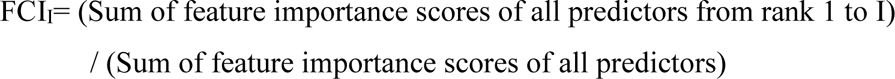

For each model optimal ‘I’ was identified the minimum value for which FCI_I_ exceeded 0.90 (accounted for 90% of the model performance). The top predictors were identified as the top I (I corresponding to this optimal value) taxa based on their feature importance scores.

Finally, for a given Bifidobacterial trait and cohort-type (16S/WGS, Infant/Adult/Senior, IndustrializedUrban/UrbanRuralMixed/RuralTribal), the consensus top informative taxa were identified as those which were identified as top predictors in at least 20% of the cohorts of that type.

### Computation of Association-Scores for the different Bifidobacterial traits

The consensus top predictors identified above were then tested for their association with the corresponding *Bifidobacterium* properties/traits. For a given cohort-type (categorized based on profiling strategy: 16S or WGS; age-group: infant or adult or senior and; life-style: IndustrializedUrban or UrbanRuralMixed or RuralTribal) and *Bifidobacterium* trait, for each taxon in the list of the corresponding top predictors, we computed the Spearman Correlation of the taxon with the *Bifidobacterium* trait (either the total number of *Bifidobacterium* or the abundance of any of the eight *Bifidobacterium* species) in each of the study-cohorts belonging to the specific cohort-type. We counted the total number of study-cohorts where the given taxa had a positive correlation and a negative correlation as SP and SN, respectively. Given the total number of study-cohorts for that specific cohort-type as T, the Association-Score for the given taxon with the specific *Bifidobacterium* trait in a given cohort-type (categorized above) as:

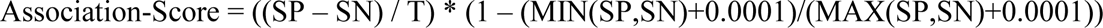

where, MAX(SP,SN) refers to the maximum of SP and SN and MIN(SP,SN) refers to the minimum of the two values.

### Computation of Association-Scores for different Disease-groups

The detailed description regarding the computation of Association-Scores of the different non-Bifidobacterial taxa with the nine different Bifidobacterial properties, including the collation of additional study-cohorts with matched diseased-control samples, selection of specific diseases for this investigation and final computation of disease-specific Association scores is described in **Text S2**.

### Description of the eight Bifidobacterium-associated intervention trials cohorts for external validation and validation of receptive-scores

Here, we collected data from eight *Bifidobacterium*-associated intervention trials (listed in **Table S13**; See **Text S3**). While one trial Bazanella *et al* was on infants/new-borns^15^, the rest seven were on adult/senior populations. Three of these cohorts were 16S sequenced^15,27,29^, and the rest were profiled using WGS^11,24–26,28^. For each of the trials, besides investigating the reproducibility of the association-patterns (identified in our analyses of the discovery cohorts described above), we also identified specific *Bifidobacterium*-associated property whose response was to be investigated in our validation of the concept of Receptive-Scores (described below and in the subsequent sections).

Five of these trials used single *Bifidobacterium* probiotics: Three using *B. animalis*^24,26,27^; one using *B. breve*^28^; one using *B. longum*^11^ and one using *B. adolescentis*^29^. For each of these trials, the target *Bifidobacterium* ‘property’ (described later) to be investigated were the abundances of the administered *Bifidobacterium* species. One of the trials used a combination of three Bifidobacteria (*B. longum*, *B. breve*, *B. bifidum*)^15^. Thus, here, there were three targeted *Bifidobacterium* properties, corresponding to the three administered Bifidobacterial taxa. One of the trials did not administer any probiotic taxa but a prebiotic (Galacto-oligosaccharide or GOS), that is expected to enrich the already resident gut *Bifidobacterium*^25^. Thus, the target property to be investigated here was the overall *Bifidobacterium* detection in a gut microbiome.

### Investigating the reproducibility of the Association-Scores

The detailed methodology for investigating the reproducibility is described in **Text S4**.

### Formulation of Receptive-Scores

For a given *Bifidobacterium* property (the property could be the abundances of any of the eight Bifidobacteria or the overall detection of *Bifidobacterium* species) and taxonomic composition profile of a microbiome of a given category (based on age-group, cohort-life-style, and microbiome-profiling-strategy), the Receptive-Score was computed as follows. We first considered the relative abundances of all non-*Bifidobacterium* taxa whose Association-Scores for the specific *Bifidobacterium* property were available (i.e. computed from our investigation of the discovery cohorts). Using these values, we first computed an Expected-Representation-Score for that property in the given microbiome as the weighted cumulated abundance of these non-Bifidobacterial taxa, where the abundances of each non-Bifidobacterial taxa was multiplied by a weight, which was simply the Association-Score obtained for that taxa with the specific *Bifidobacterium* property in microbiomes of the same category (as described above). For a matrix with multiple microbiome profiles from a specific cohort, this was performed using an inner multiplication of this abundance profile matrix with the corresponding Association-Score vector (containing the Association-Scores for the various non-Bifidobacterial taxa arranged in the same order as the columns of the abundance profile matrix). This Expected-Representation-Score for each microbiome was then ‘rank-scaled’ to values between 0 and 1 by considering all the baseline microbiomes of a cohort as below. For a given array of values ‘x’ of length k, denoted by x = {x_1_, x_2_ .., x_k_}, the values are first ranked. Next, the corresponding rank-scaled values, denoted by x_R_ = {x_1R_, x_2R_.., x_kR_}, of this set were obtained as:

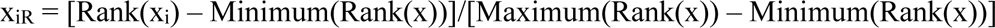

The transformation was essentially an approach of ranking the values and then converting the ranks to values between 0 and 1. We also applied the same rank-scale functions on the baseline values of the target *Bifidobacterium* property (i.e. Actual-Ranked-Baseline-Score). The Receptive-Score for the *Bifidobacterium* property was then computed as:

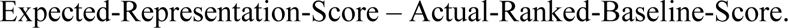

### Correlating baseline Receptive-Score, ranked baseline abundance with post-treatment increase

For each of the eight trials, we identified the specific Bifidobacterium-associated property to be investigated. For the three trials involving *B. animalis*, the target property was the abundance of *B. animalis*. For the *B. longum* supplementation Maldonado-Gomez *et al* trial and the *B. breve* supplementation trial of Grønbæk *et al*, the target properties were *B. longum* and *B. breve* abundance, respectively. Since the trial by Looijestein *et al* involved the administration of GOS that promotes the growth of all *Bifidobacterium*, the target property chosen here was the overall *Bifidobacterium* detection. For the Bazanella *et al* trial that involved supplementation of a consortia of three *Bifidobacterium* taxa (*longum*/*breve*/*bifidum*), there were three target properties, namely the abundances of *B. longum*, *B. breve* and *B. bifidum*. Another trial involving supplementation of *B. adolescentis* by Ramakrishnan *et al* was considered, where the target property was abundance of *B. adolescentis*.

The objective here was to investigate whether the Receptive-Score was significantly associated with post-treatment increase of the corresponding target properties. Since both Receptive-Score as well as the ranked post-treatment increase were calculated with respect to ranked baseline values of the target *Bifidobacterium* property, it was important to adjust for this confounding effect as it may lead to spurious correlations. We did that using robust linear regression models as:

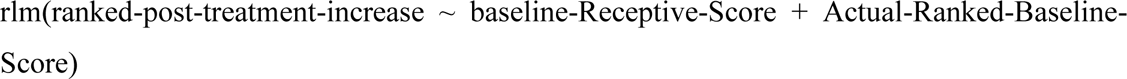

The ranked-post-treatment-increase was simply the differences between the rank-scaled abundances at the immediate post-treatment time-point and the baseline (computed using the rank-scale function described above). The robust linear regression (RLR) models were built using the rlm function of the MASS package (v 7.3.60) of R.

The RLR model was as follows:

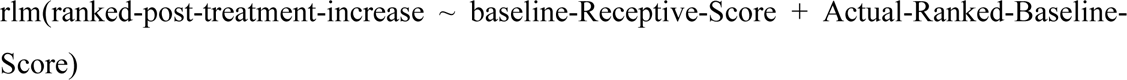

Here, the predictors were the baseline-Receptive-Score and the Actual-Ranked-Baseline-Score.

### Cross-cohort validation in B. animalis-associated trials

In the cross-cohort validation, for each of the *B. animalis*-associated trials, we trained linear-regression models to predict ranked-post-treatment-increase using the baseline-Receptive-score and the Actual-Ranked-Baseline-Score (i.e the baseline *B. animalis* abundance rank-scaled) on that cohort and test the same on the remaining two cohorts.

### Curation of a high-quality genomes from different databases and their genome annotation

We systematically probed multiple publicly available databases of reference genome and metagenome-assembled genomes to create a collation of 28,991 genomes belonging to 72 non-Bifidobacterial taxa for which which Association-Scores were obtained against multiple Bifidobacterial traits (**Figure 3B**; **Figure S19**). **Text S5** describes the detailed methodology adopted for this purpose.

### Mean Functional Abundance Profile and Conservation Profile of unique non-Bifidobacterial species-level taxa and grouping based on functional redundancy

The genome-level functional profiles (28,991 genomes X 36917 functional features) were then aggregated to obtain species-taxa-level functional features (72 taxa X 36917 functional features) using two approaches. We computed the Mean Functional Abundance profile by computing the mean abundance of each functional feature across all genomes of a given species to obtain a 72 species-level-taxa X 36917 functional feature profile matrix. Similarly, a Conservation profile (72 species-level-taxa X 36917 functional features) was computed by obtaining the detection percentage of each functional feature across all genomes of each of the 72 species-level-taxa.

Subsequently, we reduced the dimensions of the Mean Functional Abundance matrix by removing functionally redundant features originating from different schema in a step-wise manner. First, we removed functional features that did not show >= 90% conservation across the genomes of at least one of the 72 taxa. Subsequently, the functional features showing zero Euclidean distance in terms of their mean abundance across all taxa were grouped using a Greedy approach, where each functional feature was sequentially compared with all other features preceding it in the profile. If any of the preceding functional-features showed exact similarity of the mean abundance profile with the given feature, there were grouped as one functional-group. Else, it was assigned its own functional-group. Finally, we obtained a set of 16,077 functional groups from 72 non-Bifidobacterial taxa.

### *Machine-learning based identification of genome-specific functional groups of different taxa that are associated with their* Association-Scores *with different Bifidobacterial taxa and overall Bifidobacterium detection*

The next goal was to investigate whether the functional-groups of non-Bifidobacterial taxa had the ability to predict the different Bifidobacteria-linked Association-Scores. For this purpose, we adopted a two-stage Random Forest (RF)-based approach. In the first stage, we trained nine ‘Initial’ models to predict the Association-Scores for each of the eight Bifidobacterial taxa and the overall *Bifidobacterium* detection based on the mean abundance of the 16,077 functional-groups as predictors. The importance-scores for the 16,077 functional-groups were obtained for each of the nine models and the functional-groups sorted based on their scores. Based on these sorted lists of functional-groups, for each of the model, we next selected the most informative features as the top set of features that cumulatively accounted for 90% of the total importance scores for the corresponding model.

In the second stage, the nine models were re-trained using these selected set of most informative features corresponding to each model. Finally, the predictive performance of each of these nine ‘Refined’ models were judged based on the correlation between the actual and the predicted Association-Scores.

### *Identifying functional features associated to different Bifidobacterial* Association-Scores

The union set of informative features for the nine models encompassed 5,730 unique functional groups. From this list, we further selected only those functional-groups that satisfied the following criteria: appearing amongst top 50 features in at least one model; present in the list of informative features of at least two of the nine models; present in >1 non-Bifidobacterial taxa; showing significant spearman correlation of its taxa-specific mean abundances with at least two Association-Scores. Finally, 129 functional-groups were obtained that satisfied all the above criteria (**Table S16)**. Procrustes analysis was performed using the procuste.randtest function in ade4 package of R (v1.7-23). Visualization was performed using ggplot2 package of R (v3.5.2).

