## Supplementary Text and Figures for "Identifying the non-Bifidobacterial hallmarks of a Bifidobacterium-receptive gut microbiome"

### Text S1

#### Determining subset of consistent gut-associated taxa for cohorts of different Sequencing Types and Age-Category

A key challenge of gut microbiomes is the presence of a large number of microbes that are atypical and allochthonous. These microbes are mostly derived from the diet and environment and not regular residents of the gut microbiome. This results in large number of sparsely taxa that tends to inflate the microbial diversity in the gut microbiome. Thus, for identifying consistent and reproducible microbial markers across study-cohorts, it is important to remove this 'noise'. We previously implemented specific protocols to eliminate sparsely detected microbial noise and identify a minimal set of consistently detected microbes that maximally represent the gut microbial community across all investigated microbiomes[1]. Here also, we adopted a similar strategy as described below to investigate the cohorts included in this investigation. The objective of this investigation was to select the smallest possible list of microbes that could cumulatively account for the maximum fraction of the studied gut microbiomes across all study-cohorts.

To identify these taxa subsets, we divided our study-cohorts into four groups, based on different combinations of age-category (infant and adult/senior) and Sequencing Types (16S and WGS). For each group, we adopted the following approach.

To get the list of taxa for a specific study-group (16S-Infant or WGS-Infant or 16S-Adult/Senior or WGS-Adult/Senior), we defined two terms or thresholds for cohorts of that group:

1. Threshold 'X': Percentage of microbiomes in a given study-cohort where a given taxa is detected
2. Threshold 'Y': Percentage of study-cohort where a given tax needs to be detected in at least X% of the microbiomes

Now the next goal was to find a list of N taxa that is in at least X% of the microbiomes in at least Y% of the study-cohorts, such that:

a. N is reasonably small
b. N taxa cumulatively represent as much of the gut microbial community composition as possible across all investigated cohorts

Subsequently, we used a grid search approach where the values of each of X and Y were varied from 0.05 to 0.95 at a difference of 0.05 (i.e. 0.05, 0.1, 0.15, etc till 0.95). Each combination of X and Y provided different sets of microbes and the sizes of these microbial sets are noted in **Figure S3**, where the sets were determined separately for cohorts of different categories. These categories were Infant-16S, Infant-WGS, Adult/Senior-16S and Adult/Senior-WGS)

The next task was to investigate how good was to determine how maximally each microbial set represented the gut microbial community across all gut microbiomes from the corresponding category of study-cohorts. This was judged by computing two scores:

- Score 1: The proportion of microbiomes where the selected species accounted for more than 90 percent of the entire microbiome community

- Score 2: The proportion of microbiomes where the selected species accounted for less than 70 percent of the entire microbiome community

Our objective was to identify for each of the four cohort categories the smallest affordable set of microbes with the highest possible Score 1 and the lowest possible Score 2. Using this logic, we investigated each microbial set for Score 1 and Score 2 and selected the following microbial taxa sets as described below. For Infant 16S cohorts, we shortlisted 164 taxa (x=0.35, y=0.15). This set cumulatively contributed to greater than 90% relative abundance in 100% of microbiomes in Infant cohorts sequenced using 16S approach (Score 1) and less than 70% in none (Score 2). For Adult/Senior 16S cohorts, we shortlisted the microbial set with 194 taxa (x=0.4, y=0.1). This set cumulatively accounted for greater than 90% relative abundance in 90.9% of the microbiomes of cohorts belonging to this category and less than 70% in none. Similarly for the Infant cohorts sequenced using WGS, we shortlisted 226 taxa (x=0.05, y=0.1), which accounted for >=90% abundance in 93.3% microbiomes in cohorts of this category and less than 70% in none. Finally, in this manner, for the Adult/Senior WGS cohorts, we shortlisted 139 taxa (x=0.35, y=0.1), which accounted for which the Score 1 was 90.3% and Score 2 was 2.03%.

### Text S2

### Investigation of the association-patterns of the non-Bifidobacterial taxa with the *Bifidobacteria* in different Diseased-groups

*Collation of datasets from different diseases and disease-matched controls*

Since a majority of the study-cohorts investigated here were from the IndustrializedUrban cohorts and from the adult/senior age-groups (with minimal datasets containing patient gut microbiomes from Non-Industrialized populations), we focused on investigating the variations of association-patterns (of different non-Bifidobacterial taxa with the different Bifidobacteria) across different diseases only in these populations. Further, to gain a comprehensive understanding of the disease-associated alterations in these association-patterns, we further supplemented our original data collection of 45,809 gut microbiomes from 121 study-cohorts with an additional 27 study-cohorts containing a total of 5,249 gut microbiomes from different diseases with matched non-diseased controls (Table S1)[1–27].

Addition of these cohorts resulted in a total of 69 study-cohorts containing gut microbiomes from nine different diseases along with the microbiomes from matched study-specific control (non-diseased) individuals from from Adult/Senior age category and IndustrializedUrban population. These nine diseases were Polyps, Colorectal Cancer (CRC), Type-II Diabetes (T2D), Inflammatory Bowel Disorder (IBD), Cardiovascular Diseases (CVD), Parkinson's Disease, COVID, Prediabetes and Liver Disease. We considered nine diseases as they contained data from more than three study-cohorts, thus increasing the reliability of the computed associations.

Out of these 69 study-cohorts thus selected, 34 and 35 datasets were from 16S and WGS profiling strategy respectively, forming a pooled set of 15,095 microbiomes which were finally utilized for disease-wise Association-Scores, thereby enabling the investigation of the replicability of the patterns across different profiling strategies (see **Table S1**)

*Computation of Association-Scores for the different Bifidobacterial traits*

The consensus top predictors identified while computing the overall Association-Scores were investigated for their association with the corresponding Bifidobacterium properties/traits. For a given disease category (it can be one of the nine diseases or the pooled controls) and Bifidobacterium trait, for each taxon in the list of the corresponding top predictors, we computed the Spearman Correlation of the taxon with the Bifidobacterium trait (either the total number of Bifidobacterium or the abundance of any of the eight Bifidobacterium species) in each of the study-cohorts belonging to the specific disease category. We counted the total number of study-cohorts where the given taxa had a positive correlation and a negative correlation as SP and SN, respectively. Given the total number of study-cohorts for that specific cohort-type as T, the Association-Score for the given taxon with the specific Bifidobacterium trait in a given disease category (categorized above) as:

$$\text{Association-Score} = ((SP - SN) / T) * (1 - (MIN(SP,SN)+0.0001)/(MAX(SP,SN)+0.0001))$$

where, MAX(SP,SN) refers to the maximum of SP and SN and MIN(SP,SN) refers to the minimum of the two values.

*Investigating the reproducibility of Disease-wise Association-Scores across different profiling strategies*

As, we are performing the analysis using datasets both from 16S and WGS profiling strategy, we investigated the reproducibility of disease-wise Association-Scores when calculated for 16S and WGS study-cohorts separately. For this analysis, we considered those disease categories only for which more than 3 datasets are available from each of the 16S and WGS sequence types and hence, we considered CRC, CVD, IBD and pooled controls for investigating the reproducibility. We calculated the Association-Scores using the same formula as described above, but for 16S and WGS study-cohorts separately, and finally we compared the Association-Scores within a particular disease category stratified by 16S and WGS profiling strategy for each *Bifidobacterium* feature using Spearman correlation (see **Figure S16**)

*Investigation of association-patterns across different diseases as well as with non-diseased state*

For each of the nine Bifidobacterium features, spearman correlation was calculated using the Association-Scores across all the nine diseases as well as controls to check if there is pattern of association among the disease groups as compared to control. These inter-disease association-patterns have been shown as heatmaps in **Figure S17**.

### Text S3

**Descriptions of Unseen Validation Datasets utilized for investigating the reproducibility of the Association-Scores**

We investigated eight datasets from different intervention trials that relate to previously published studies (detailed in **Table S13**) to confirm the patterns of Association-Scores as a validation.

Below is a summary of the datasets and the different analyses conducted on the datasets.

1. ***Maldonado-Gomez_2016***: This study was a double-blind, placebo-controlled crossover trial performed in the United States, including 23 healthy adult participants aged from 22 to 38 years[1]. Participants followed an industrialized-urban lifestyle and received a daily dose of *Bifidobacterium longum* subsp. *longum* AH1206 during two separate 7-week testing phases, each consisting of a baseline, a 14-day treatment period, and a 28-day follow-up. Whole-genome shotgun sequencing (WGS) of stool microbiomes was performed and the taxonomic profiling was done using Metaphlan 3[2] to get the species-level taxa abundance profiles for the microbiomes and for our analysis, we utilized data for 21 subjects (baseline), as the metadata was available for 21 subjects only including the baseline. A notable aspect of the study was the stratification of subjects into "persisters" and "non-persisters" based on whether the administered *B. longum* strain established long-term colonization. Prior analyses in the study revealed that baseline microbiome features particularly the abundance of carbohydrate metabolism genes, differentiated these two groups, suggesting that functional traits of the gut microbiome influence probiotic engraftment potential.
   For the above-mentioned groups (persisters and non-persisters), we applied our 'Receptive-Score' (see Methods) methodology to the baseline WGS profiles of the 21 individuals. Using *B. longum*-specific Association-Scores trained in industrialized adult cohorts, we computed Receptive-Scores and tested their ability to distinguish persisters from non-persisters. Indeed, baseline Receptive-Scores were significantly higher among persisters (Mann-Whitney U-test **P = 0.014**), confirming that our framework has the potential to identify individuals with greater probiotic receptivity. Additionally, this dataset was used also included in evaluating whether the association-patterns identified in the discovery cohorts could be reliably reproduced in the validation cohorts.

2. ***ZhangQ_2024***: This study was a randomized, parallel-group, placebo-controlled clinical trial carried out in China involving 200 adult individuals from IndustrializedUrban lifestyle, with gut microbiomes profiled using whole-genome shotgun (WGS) sequencing[3]. The taxonomic profiling was done using Metaphlan 3[2] to get the species-level taxa abundance profiles for the microbiomes. The study investigated the impact of *Bifidobacterium animalis* subsp. *lactis* BL-99 supplementation. Participants first underwent a 2-week run-in phase with dietary restrictions on probiotic-containing foods, followed by 8 weeks of treatment (BL-99, PPI, or placebo) and an 8-week follow-up. Fecal samples were collected, of which we utilized 350 microbiomes from 175 subjects at baseline and post-treatment.

This dataset was utilized in several validation analyses. First, we utilized the dataset in evaluating whether the association-patterns identified in the discovery cohorts could be reliably reproduced in the validation cohorts. Second, we demonstrated that microbiome-level Receptive-Scores, computed using Association-Scores and baseline abundances of non-*Bifidobacterium* taxa, significantly predicted the post-treatment increase of *B. animalis* using Leave-One-Out-Cross-Validation (**Coeff = 0.23, P = 0.03**) and robust linear regression model (**Coeff = 0.31, P = 0.007**) (see Methods). Third, the dataset was used to assess the robustness of the Receptive-Score framework through a cross-cohort validation approach: a cohort-specific robust linear regression model was trained to predict *B. animalis* increase based on baseline Receptive-Score and baseline abundance, and then tested on other intervention cohorts. This approach confirmed the generalizability and predictive power of the Receptive-Score methodology across independent datasets.

3. ***LooijesteijnE_2024***: This study was a double-blind, randomized, parallel-group clinical trial carried out in the Netherlands involving 86 healthy women aged 40 to70 years, from IndustrializedUrban lifestyle[4]. The intervention tested the effect of galacto-oligosaccharides (GOS) supplementation at two doses over three weeks. After a control period lasting three weeks, participants were divided based on their dietary fiber consumption, body mass index (BMI), and age, and then randomly assigned to different treatment groups. Stool samples were gathered at three different intervals: at the beginning of the control phase (t = −3), at the conclusion of the control phase and the commencement of treatment (t = 0), and at the end of the treatment period (t = 3). For our analysis, we considered the end of the control period (t = 0) as baseline and post-treatment (t = 3) as the outcome timepoint, resulting in 258 microbiomes from 86 subjects. The study mainly concentrated on changes in fecal Bifidobacterium levels and the composition of gut microbiota, evaluated through whole-genome shotgun (WGS) sequencing. The taxonomic profiling was done using Metaphlan 3[2] to get the species-level taxa abundance profiles for the microbiomes.
   This dataset was utilized in several validation analyses. First, we utilized the dataset in evaluating whether the association-patterns identified in the discovery cohorts could be reliably reproduced in the validation cohorts. Second, we demonstrated that microbiome-level Receptive-Scores, computed using Association-Scores and baseline abundances of non-*Bifidobacterium* taxa, significantly predicted the post-treatment increase of *Bifidobacterium* using Leave-One-Out-Cross-Validation (**Coeff = 0.30, P = 0.005**) and robust linear regression model (**Coeff = 0.17, P = 0.01**).

4. ***SunB_2022***: This study was a randomized, double-blind, placebo-controlled clinical trial carried out in China to examine the effects of *Bifidobacterium animalis* subsp. *lactis* Probio-M8 on patients with coronary artery disease (CAD)[5]. A total of 71 senior participants were randomized to receive either the probiotic (37 subjects) or a placebo (34 subjects) over a six-month period, during which the probiotic group was administered daily sachets of Probio-M8. Fecal samples were collected from a subset of 41 individuals (24 from the probiotic group and 17 from the placebo group) at three time points: baseline (day 0), midpoint (day 90), and end of treatment (day 180). We focused on the 24 probiotic-group individuals, contributing 72 microbiomes profiled using whole-

genome shotgun (WGS) sequencing and the taxonomic profiling was done using Metaphlan 3[2] to get the species-level taxa abundance profiles for the microbiomes. The study aimed to assess microbiota and metabolic changes, especially in the context of adjunctive probiotic therapy for CAD.

This dataset was utilized in several validation analyses. First, we utilized the dataset in evaluating whether the association-patterns identified in the discovery cohorts could be reliably reproduced in the validation cohorts. Second, we demonstrated that microbiome-level Receptive-Scores, computed using Association-Scores and baseline abundances of non-*Bifidobacterium* taxa, predicted the post-treatment increase of *B. animalis* at 90 days (LOOCV: **Coeff = 0.5, P = 0.03**; RLR: **Coeff = 0.7, P = 0.009**) and at 180 days (LOOCV: Coeff = 0.45, P = 0.06; RLR: Coeff = 0.2, P = 0.36). The reduced statistical significance recorded at 180 days probably indicate a decrease in the probiotic impact over time, which aligns with the gradual reduction in the abundance of B. animalis after treatment, implying that there is limited long-term retention after stopping the intervention. Third, the dataset was used to assess the robustness of the Receptive-Score framework through a cross-cohort validation approach: a cohort-specific robust linear regression model was trained to predict *B. animalis* increase based on baseline Receptive-Score and baseline abundance, and then tested on other intervention cohorts. This approach confirmed the generalizability and predictive power of the Receptive-Score methodology across independent datasets.

5. ***BazanellaM_2017***: This study was a double-blind randomized, placebo-controlled trial involving 106 healthy infants in Germany (IndustrializedUrban cohort), where participants were allocated to receive bifidobacteria-supplemented formula, placebo formula, or exclusive breastfeeding during their first year of life[6]. Microbiome profiling was conducted using 16S rRNA sequencing and the taxonomic profiling was done using Spingo[7] to get the species-level taxa abundance profiles for the microbiomes. From this dataset, we selected matched subjects with samples available at 7, 12, and 24 months, resulting in 87 subjects and 173 microbiome samples.

   This dataset was utilized in several validation analyses. First, we utilized the dataset in computing the associations between the relative abundances of non-*Bifidobacterium* taxa and each of seven *Bifidobacterium* species (excluding *B. animalis*). These association-patterns were then compared to those derived from discovery cohorts within the same study category (Infant–IndustrializedUrban–16S). We observed significant positive correlations (P-value > 0.05) for these association-patterns (spearman coefficient ranging from 0.27 to 0.55), with 4 out of 7 species showing correlations ≥ 0.40, indicating robust cross-cohort reproducibility. Second, we demonstrated that microbiome-level Receptive-Scores, computed using Association-Scores and baseline abundances of non-*Bifidobacterium* taxa, predicted the increase of *B. longum, B. bifidum, B. breve* at 7 months, 12 months and 24 months. The significant increase of *B. breve* was predicted at 7 months (LOOCV: **Coeff = 0.6, P = 1.00E-05**; RLR: **Coeff = 0.5, P = 0.003**) whereas for *B. bifidum* it was predicted at 7 (LOOCV: **Coeff = 0.59, P = 2.00E-05**; RLR: **Coeff = 0.41, P = 0.03**) as well as at 12 months (LOOCV: **Coeff = 0.6, P = 1.00E-05**; RLR: **Coeff = 0.37, P = 0.05)**. Predictive strength generally remained high at 12 and 24 months for all three species, the probable reason might be the increased variability in long-term response.

6. ***BaZ_2021***: This study was a randomized, four-period, crossover model conducted in a partially blinded fashion with adult participants from USA (IndustrializedUrban lifestyle)[8]. Participants received four treatments in randomized order: (A) yogurt smoothie without BB-12 (YS), (B) yogurt smoothie with BB-12 added post-fermentation (POST), (C) yogurt smoothie with BB-12 added pre-fermentation (PRE), and (D) BB-12 capsule (CAP), with each treatment lasting 4 weeks and a 2-week washout period in between. Measurements included anthropometrics, biochemical markers, physical activity, and immune endpoints. Microbiome profiling was conducted using 16S rRNA sequencing and the taxonomic profiling was done using Spingo[7] to get the species-level taxa abundance profiles for the microbiomes. For our analysis, we took 24 matched subjects for baseline, PRE, POST and CAP covering 111 microbiomes. This dataset was utilized in several validation analyses. First, we demonstrated that microbiome-level Receptive-Scores, computed using Association-Scores and baseline abundances of non-*Bifidobacterium* taxa, significantly predicted the increase of *B. animalis* at PRE (LOOCV: **Coeff = 0.73, P = 0.0003**; RLR: **Coeff = 0.84, P = 0.003**), at POST (LOOCV: **Coeff = 0.8, P = 0.00013**; RLR: **Coeff = 0.58, P = 0.068**), and at CAP (LOOCV: **Coeff = 0.76, P = 0.00017**; RLR: **Coeff = 0.71, P = 0.03**). Second, the dataset was used to assess the robustness of the Receptive-Score framework through a cross-cohort validation approach: a cohort-specific robust linear regression model was trained to predict *B. animalis* increase based on baseline Receptive-Score and baseline abundance, and then tested on other intervention cohorts. This approach confirmed the generalizability and predictive power of the Receptive-Score methodology across independent datasets.

7. ***GronbaekI_2025***: This study was a randomized, double-blind, placebo-controlled investigation[9]. Participants were given either Bif195 or placebo capsules each day for an 8-week period, followed by another 8 weeks without treatment. Stool samples were gathered during every visit for microbiome evaluation, along with surveys measuring disease severity and quality of life. This dataset employed WGS sequencing data from adult participants in Denmark (DNK), belonging to an IndustrializedUrban lifestyle cohort. The taxonomic profiling was done using Metaphlan 3[2] to get the species-level taxa abundance profiles for the microbiomes. The therapeutic agent used was *Bifidobacterium breve*. We took a total of 14 matched subjects encompassing 56 microbiomes across four time-points: Baseline, Week 4, Week 8, and Week 16.
   This dataset was utilized in several validation analyses. First, we utilized the dataset in evaluating whether the association-patterns identified in the discovery cohorts could be reliably reproduced in the validation cohorts. Second, we demonstrated that microbiome-level Receptive-Scores, computed using Association-Scores and baseline abundances of non-*Bifidobacterium* taxa, significantly predicted the increase of *B. breve* at Week 4 (LOOCV: **Coeff = 0.69, P = 0.01**; RLR: **Coeff = 0.69, P = 0.03**). However, the predictions for *B. breve* increase at Week 8 and Week 16 were not statistically significant using baseline. We also used Week 8 as the baseline, since it marked the completion of the intervention and computed Receptive-Scores using Association-Scores and Week 8 abundances of non-*Bifidobacterium* taxa. This predicted the post-treatment increase of *B. breve* at Week 16 (LOOCV: Coeff = 0.1, *P* = 0.76; RLR: **Coeff = 0.45, *P* = 0.001**).

8. ***RamakrishnanM_2025***: This study was a randomized, placebo-controlled trial examining the effectiveness of *B. adolescentis* VS-1 in alleviating symptoms of lactose intolerance in confirmed lactose maldigesters[10]. Participants were screened using a hydrogen breath test and randomized to either probiotic (11 subjects) or placebo (10 subjects). Those in the treatment group received *B. adolescentis* iVS-1 supplementation over the intervention period, with stool samples collected pre- and post-treatment for microbiome profiling. This dataset includes 16S sequencing data from adult participants in the USA, categorized under the IndustrializedUrban lifestyle cohort. The taxonomic profiling was done using Spingo[7] to get the species-level taxa abundance profiles for the microbiomes. We used data from 11 subjects who took probiotics in the treatment group, resulting a total of 22 microbiomes across two time-points. Using this dataset, we demonstrated that microbiome-level Receptive-Scores, computed using Association-Scores and pre-treatment abundances of non-*Bifidobacterium* taxa, predicted the post-treatment increase of *B. adolescentis* (LOOCV: Coeff = 0.527, P = 0.1; RLR: **Coeff = 0.38, P = 0.01**). Since the probiotic intervention was ongoing until the second time point, we treated the first (pre-treatment) time point as the effective baseline, as it reflects the point when the treatment was completed. Although the LOOCV outcome did not reach only marginal statistical significance, this was probably due to the limited statistical power associated with the small sample size ($n$ = 11).

### Text S4

### Detailed methodology for investigating the reproducibility of the Association-Scores in the Validation Intervention Cohorts

Five of the validation intervention trial cohorts contained microbiomes of a specific type, namely from adult/senior individuals from IndustrializedUrban populations and sampled using WGS approach[1–5]. This microbiome group also dominated our discovery cohorts. Thus, first, we investigated here whether the Association-Scores computed from our discovery-cohorts showed similar patterns in this validation cohorts. Thus, we recomputed the Association-Scores of each of the non-Bifidobacterial taxa with respect to each of the nine Bifidobacterium properties (as described above) in specifically these five validation cohorts and subsequently correlated these values with the original Association-Scores obtained for each of the corresponding nine properties in the discovery cohorts. Since the five validation cohorts corresponded to adult/senior individuals from IndustrializedUrban with WGS profiling approach, for the correlation analysis, we took the mean of the original Association-Scores computed form adult and senior individuals, from IndustrializedUrban populations using WGS.

Only one of our validation cohorts was from Infants (IndustrializedUrban population using 16S profiling)[6]. Thus, to investigate the reproducibility of the Infant-associated association-patterns, we first computed the spearman correlations of each of the non-Bifidobacterial taxa with eight Bifidobacterium-associated properties (with the exception of B. animalis as it did not yield any consistent pattern of associations in the discovery cohort). We then correlated this Spearman correlation patterns with the corresponding Association-Scores for each non-Bifidobacterium taxa (detected commonly across both datasets) with each of the eight Bifidobacterium properties computed for the 16S-sequenced microbiomes from Infants from IndustrializedUrban populations.

**Text S5**

**Database of reference genomes and metagenome-assembled genomes for the functional investigation**

*Creation of the database*

We collated a genome collection of 60,573 high quality genomes (with completeness > 90% and contamination < 5%), comprising of reference genomes and metagenome assembled genomes (MAGs) from adult/senior age-group as most of the MAGs were available for this age-group only, encompassing 1671 different human microbiome associated taxa. The genomes were collated from two major genome repositories: NCBI-RefSeq database containing 26,026 reference genomes and 34547 MAGs from two different repositories[1,2]. We retained 28,991 genomes belonging to 72 consistently occurring non-Bifidobacterial taxa for and finally these 28,991 genomes were utilized for generating the functional profiles for 72 non-Bifidobacterial taxa.

*Genome annotation*

All selected high-quality genomes were functionally annotated using the eggNOG-mapper pipeline (v2), which utilizes the eggNOG database (version 5.0)[3,4]. Protein-coding genes were predicted with Prodigal, integrated within the eggNOG-mapper workflow[3]. The resulting annotation files were processed using an R script to construct functional profile matrices by quantifying the occurrence of each functional feature within a genome. These matrices quantitatively capture the presence of functional signatures across genomes. Functional profiles were generated for nine different functional schemas resulting in a total of 36,917 functional features (Features break-up: CAZy: 95, KEGG Module: 619, KEGG Ortholog: 7009, KEGG Reactions: 2834, KEGG RClass: 1098, BiGG: 9468, EC Number: 2250, PFAMs: 7403, COG: 6144)[5–9].

**Text S6**

**Linear Regression Model based approach to *Association of the detection pattern of different Bifidobacteria with the cohort Sequencing-Type, Cohort-Type and host age-group categories***

For a given *Bifidobacterium* species 'i', the first linear regression models were implemented as:

*Model 1: lm (Prevalence ~ SequencingType + AgeCategory + CohortType)*

This model utilized the all the study-cohorts, where in Prevalence indicated the prevalence of the species 'i' in a given study-cohort which was regressed using the SequencingType, AgeCategory and CohortType of that study-cohort. This model computed the difference in the prevalence of species 'i' in the infant and senior age-groups with respect to the adult age-group (after adjusting for CohortType and SequencingType); the difference in the prevalence of 'i' in the RuralTribal and UrbanRuralMixed cohorts with respect to the IndustrializedUrban cohort (after adjusting for AgeCategory and SequencingType) and; the prevalence difference of 'i' in the 16S with respect to the WGS (after adjusting for AgeCategory and CohortType). The second model:

*Model 2: lm (Prevalence ~ SequencingType + AgeCategory)*

This model utilized only the infant and senior cohorts belonging to the IndustrializedUrban and computed the prevalence variation of the species 'i' only between the Infant and Senior cohorts (after adjusting for the SequencingType). We did not consider RuralMixed and UrbanRuralMixed cohorts here as most cohorts from Infants and Seniors belonged to the IndustrializedUrban category.

*Model 3: lm (Prevalence ~ SequencingType + Cohort)*

This model utilized only the RuralTribal and UrbanRuralMixed cohorts, belonging to the adult AgeCategory, and profiled the difference only between the RuralTribal and UrbanRuralMixed after adjusting for the SequencingType.

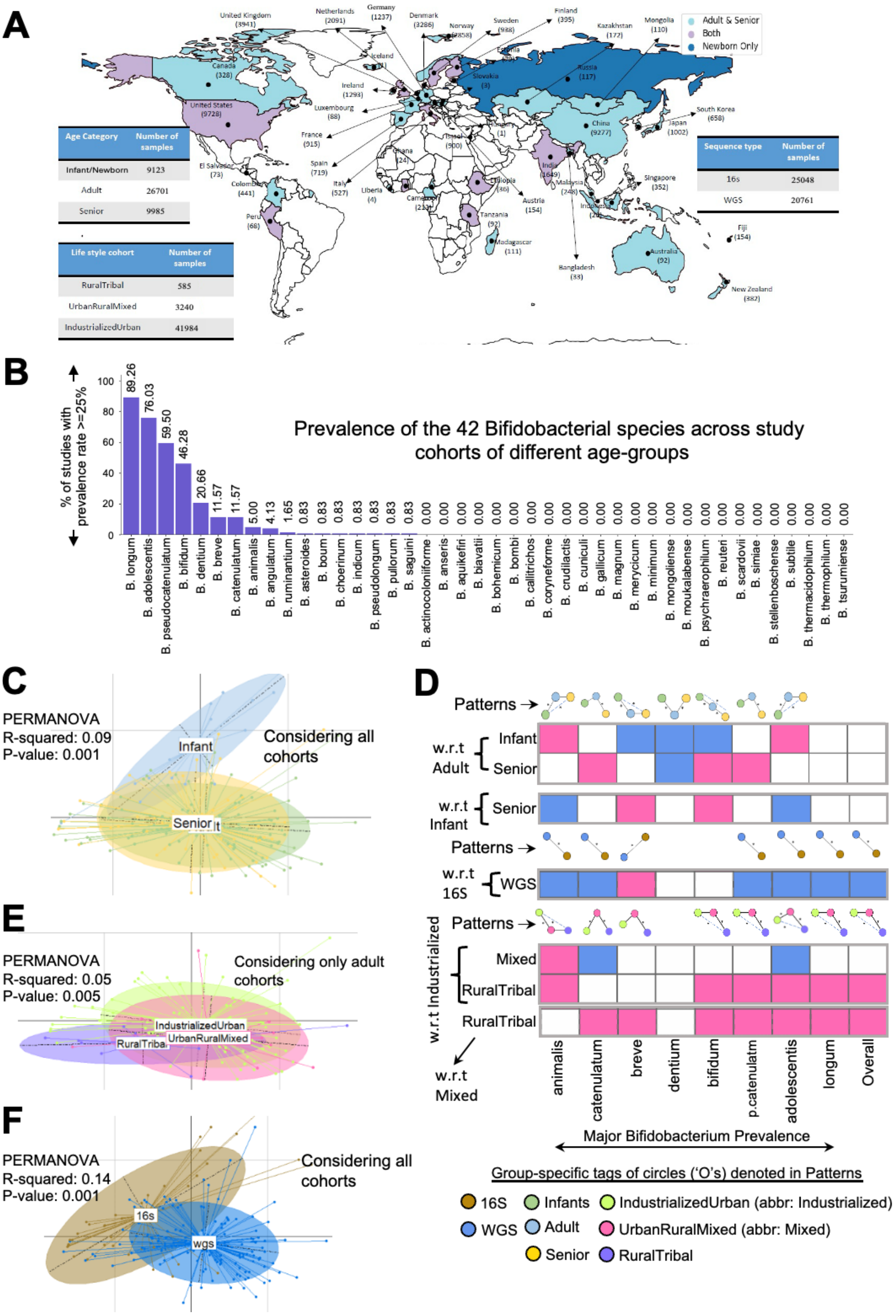


**Figure S1. A.** Geographical distribution of study-cohorts used in this investigation is illustrated on a world map. Each country is highlighted in a distinct color, representing the different age groups for which gut microbiome data were available. The total number of microbiome samples from each country is also indicated on the map. The total number of microbiomes from the different age-groups are also indicated. Also shown are the number of 16S and WGS gut microbiomes and the number of microbiomes from different Cohort-Types (IndustrializedUrban, UrbanRuralMixed, RuralTribal). **B.** Barplots showing the percentage of studies in which each of the 42 *Bifidobacterium* species reached a prevalence of >=0.25. Bars are ordered in decreasing order of study coverage. Species detected in >=0.25 prevalence in at least 5% of cohorts were considered as consistently represented and selected for further analyses. **C.** Principal Coordinate Analysis plot showing the variation in the overall prevalence pattern of the major Bifidobacterial species between the infant, adult and senior study-cohorts (considering all cohorts). Each point indicates a study-cohort. The results of the PERMANOVA analysis (R-squared, P-value) highlighting the significance of these variations is also indicated. **D.** Heatmap summarizing the results of linear regression models examining variation in the prevalence of the eight major *Bifidobacterium* species, as well as overall *Bifidobacterium* detection, across different pairwise comparisons. These include pairwise comparisons between different age groups, Cohort-Types, and Sequencing-Types, highlighting differences in species-specific and overall *Bifidobacterium* prevalence patterns. Please note comparisons between any two groups (derived from a specific study-cohort specific metadata, either, age-category or Sequencing-Type or Cohort-Type) is performed after adjusting for the remaining two metadata. For each of the three metadata, the overall patterns for each individual species (or overall detection) are also indicated. **E.** Principal Coordinate Analysis plot showing the variation in the overall prevalence pattern of the major Bifidobacterial species between the adult cohorts belonging to the different Cohort-Types (IndustrializedUrban, UrbanRuralMixed and RuralTribal). Each point indicates a study-cohort. The results of the PERMANOVA analysis (R-squared, P-value) highlighting the significance of these variations are also indicated. **F.** Principal Coordinate Analysis plot showing the variation in the overall prevalence pattern of the major Bifidobacterial species between the adult cohorts belonging to the different Sequencing-Types (16S or WGS). Each point indicates a study-cohort. The results of the PERMANOVA analysis (R-squared, P-value) highlighting the significance of these variations are also indicated.

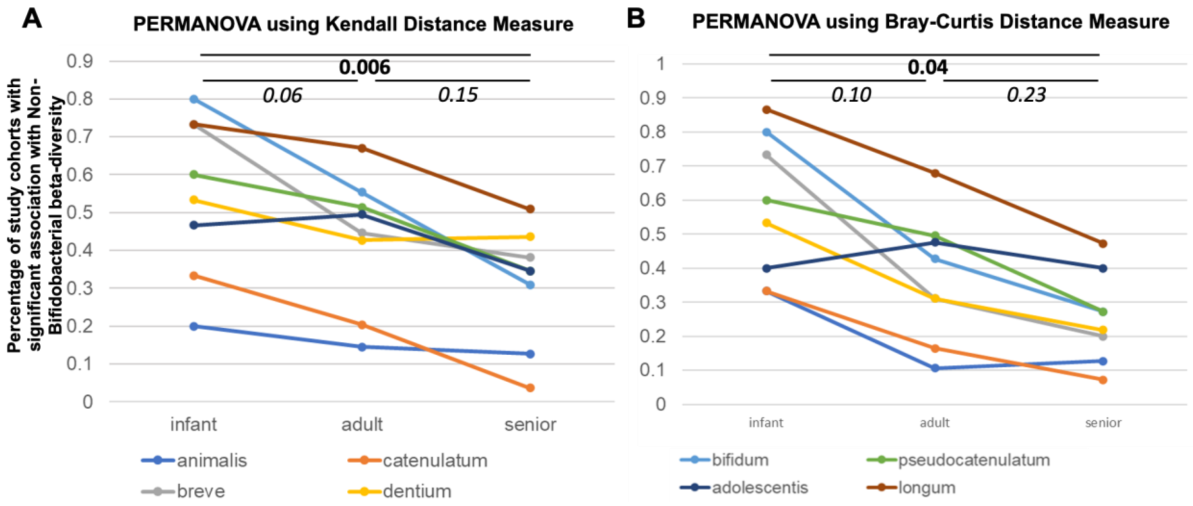


**Figure S2**. Line plots showing for each of the eight major *Bifidobacterium* species the percentage of study-cohorts with significant association with non-Bifidobacterial community compositions, when the community variations are processed using the **A.** Bray-Curtis and **B.** Kendall distance measures. The results of the Mann-Whitney tests comparing these patterns across cohorts from each age-group pair is also indicated.

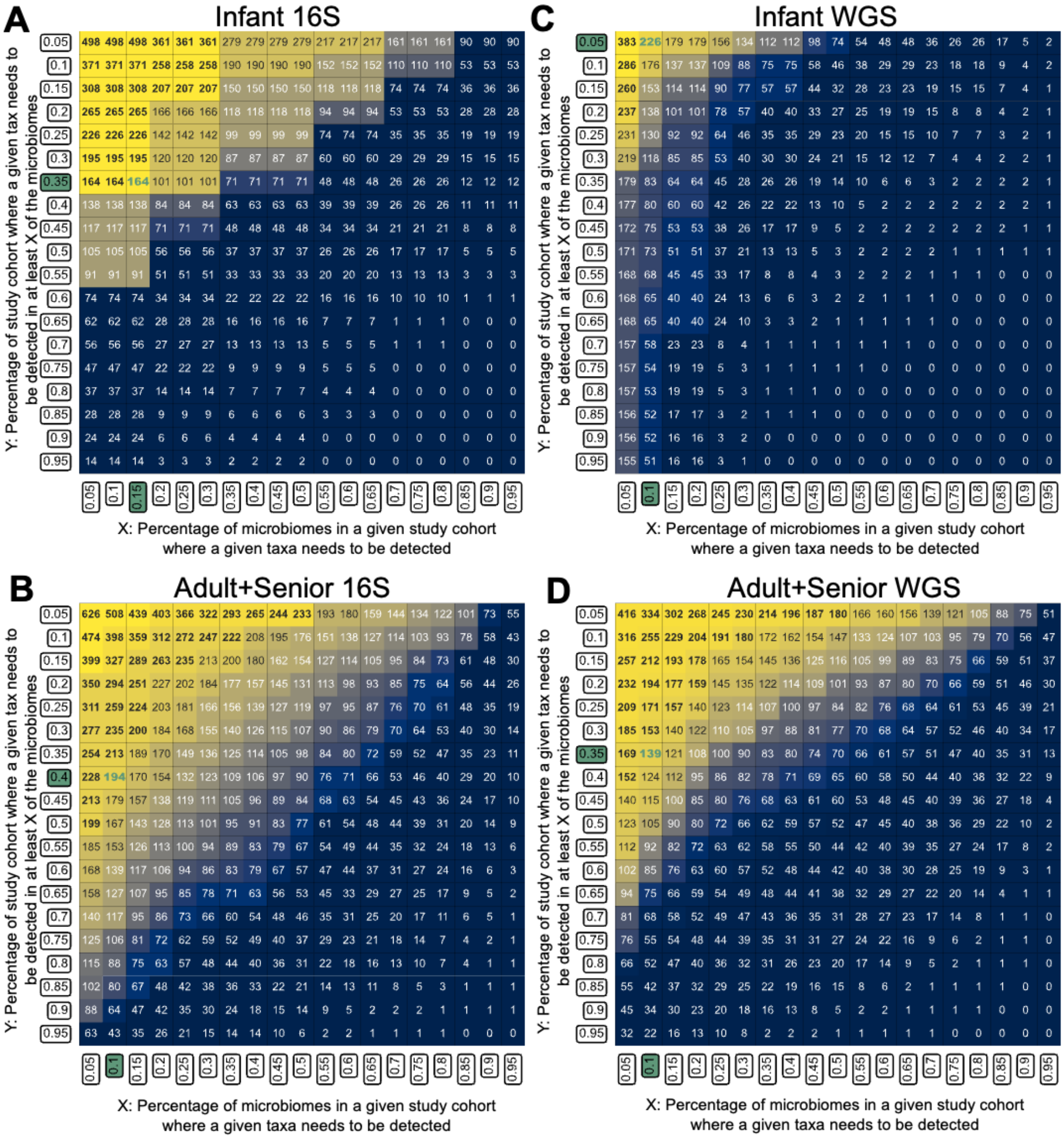


**Figure S3.** Number of taxa identified using various thresholds of X and Y, where X is the percentage of microbiomes of a given study-cohort where the taxa is detected and Y is the number of study-cohorts where the taxa is detected in at least X% of the microbiomes. These values were separately obtained for **A.** Infant-16S cohorts, **B.** Adult+Senior-16S cohorts, **C.** Infant-WGS cohorts and **D.** Adult+Senior-WGS cohorts.

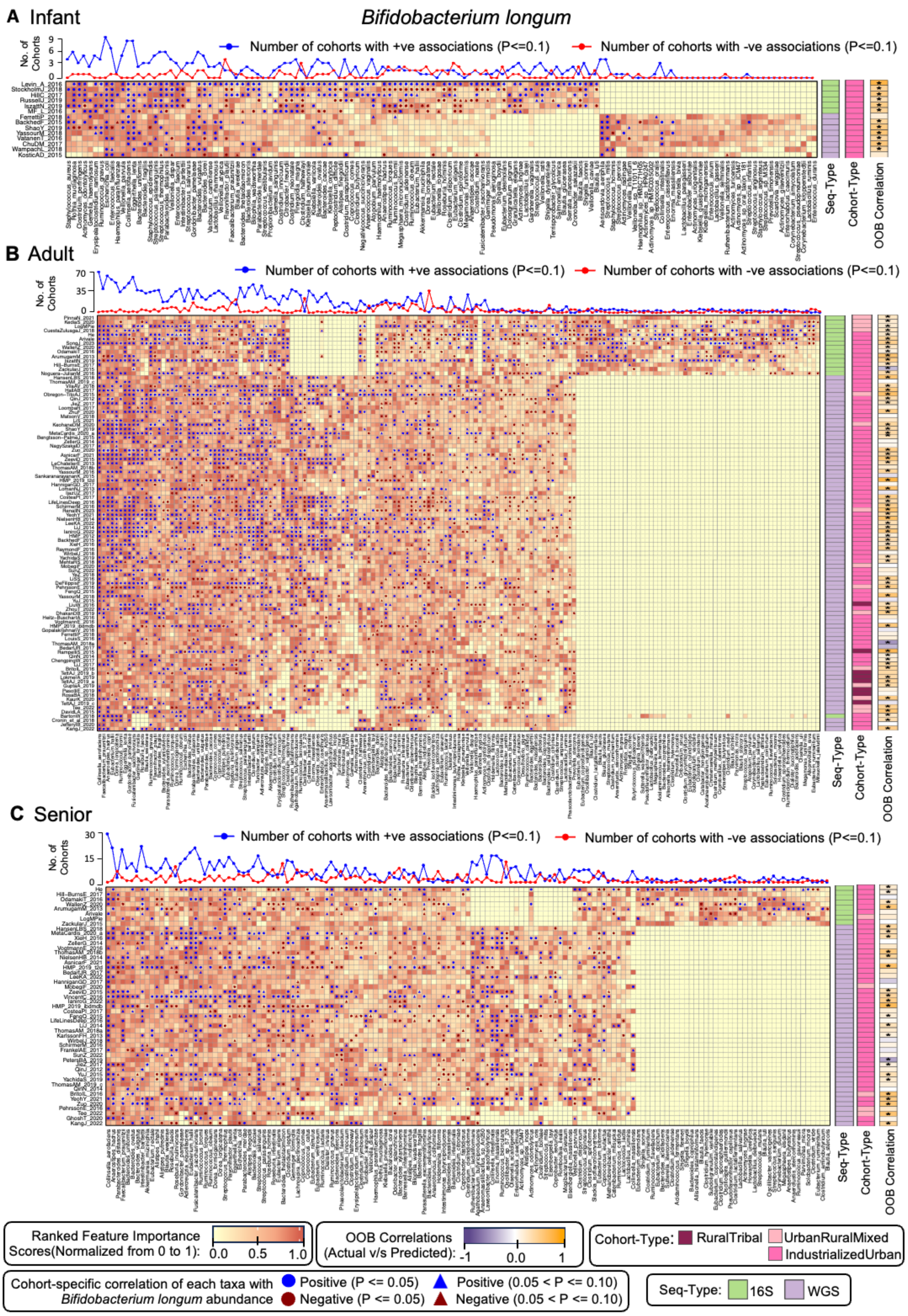


**Figure S4. Random Forest (RF) model-based identification of distinct sets of non-Bifidobacterial taxa that consistently associated with the abundance of *Bifidobacterium longum*, serving as top predictors of *B. longum* abundance in the gut microbiome. A-C.**

Heatmap showing the variation of the rank-normalized feature importance scores of the top predictor taxa for the Random Forest models built for Infant (A), Adult (B) and Senior (C) cohorts. The profiling strategy (Seq-Type), and the life-style (Cohort-Type) of each cohort is indicated as strips as shown for each heatmap. Also indicated are the out-of-bag (OOB) correlations between the actual and RF-predicted *B. longum* for each cohort. Symbols within the heatmap cells indicate both the direction (positive or negative) and strength (based on p-values) of the association between each non-Bifidobacterial taxon (columns) and *B. longum* abundance within each cohort (rows). Line plots above each heatmap summarize these associations and show the number of cohorts where each non-Bifidobacterial taxon displays significant positive or negative associations with *B. longum* abundance ($p \leq 0.1$). The extent and consistency of these associations varied by the specific non-Bifidobacterial taxon, with some showing notably higher predictive power and consistent directionality. Association patterns were also cohort-type-dependent, namely Infant cohorts exhibited distinct sets of associated taxa compared to other age groups (same for other age-groups); 16S and WGS cohorts showed differing patterns. We also observed variations depending upon the life-style of cohorts. These patterns were captured in the Association-Scores as described in **Figure 3**.

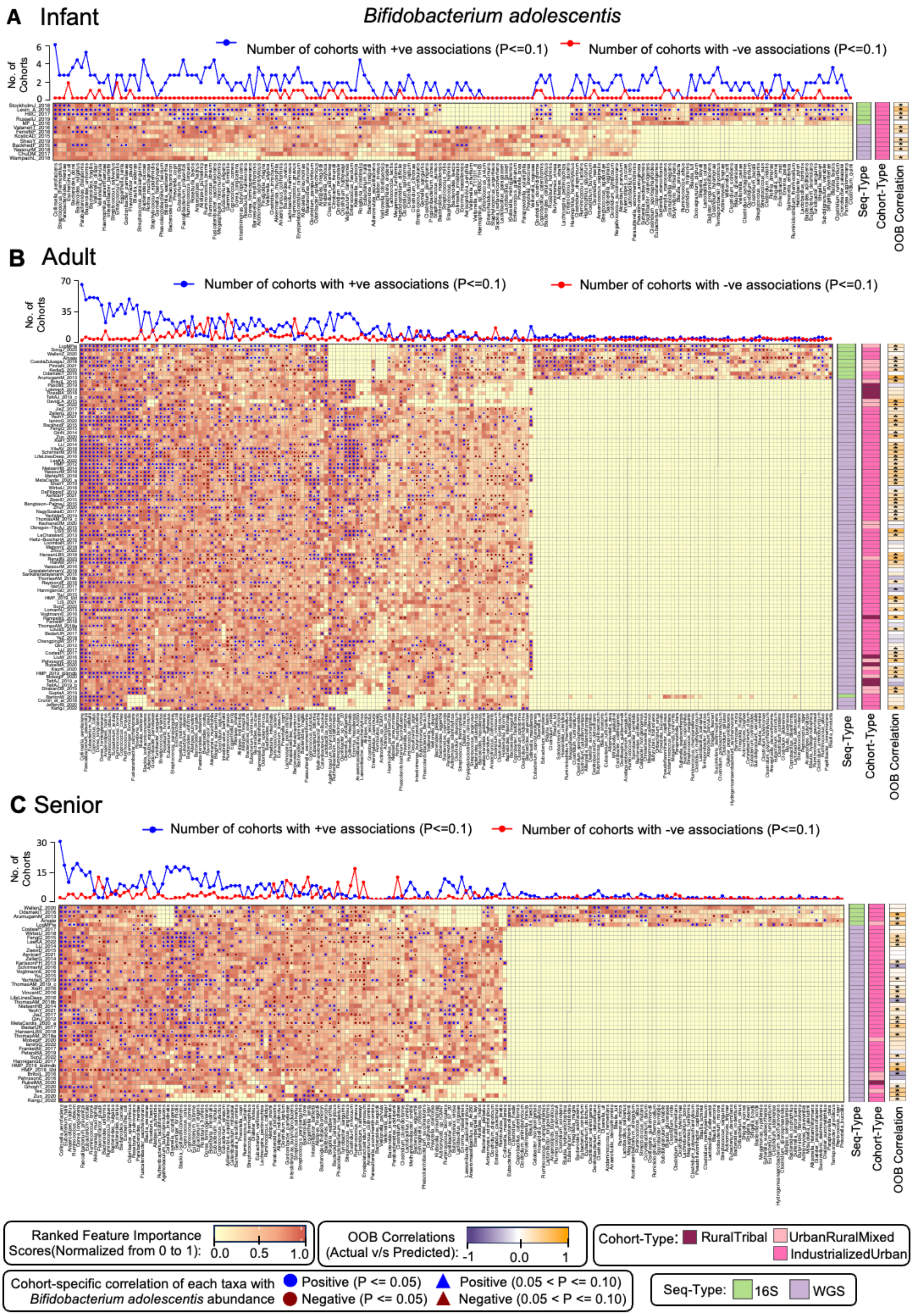


**Figure S5. Random Forest (RF) model-based identification of distinct sets of non-Bifidobacterial taxa that consistently associated with the abundance of *Bifidobacterium adolescentis*, serving as top predictors of *B. adolescentis* abundance in the gut**

**microbiome. A-C.** Heatmap showing the variation of the rank-normalized feature importance scores of the top predictor taxa for the Random Forest models built for Infant (A), Adult (B) and Senior (C) cohorts. The profiling strategy (Seq-Type), and the life-style (Cohort-Type) of each cohort is indicated as strips as shown for each heatmap. Also indicated are the out-of-bag (OOB) correlations between the actual and RF-predicted *B. adolescentis* for each cohort. Symbols within the heatmap cells indicate both the direction (positive or negative) and strength (based on p-values) of the association between each non-Bifidobacterial taxon (columns) and *B. adolescentis* abundance within each cohort (rows). Line plots above each heatmap summarize these associations and show the number of cohorts where each non-Bifidobacterial taxon displays significant positive or negative associations with *B. adolescentis* abundance ($p \leq 0.1$). The extent and consistency of these associations varied by the specific non-Bifidobacterial taxon, with some showing notably higher predictive power and consistent directionality. Association patterns were also cohort-type-dependent, namely Infant cohorts exhibited distinct sets of associated taxa compared to other age groups (same for other age-groups); 16S and WGS cohorts showed differing patterns. We also observed variations depending upon the life-style of cohorts. These patterns were captured in the Association-Scores as described in **Figure 3**.

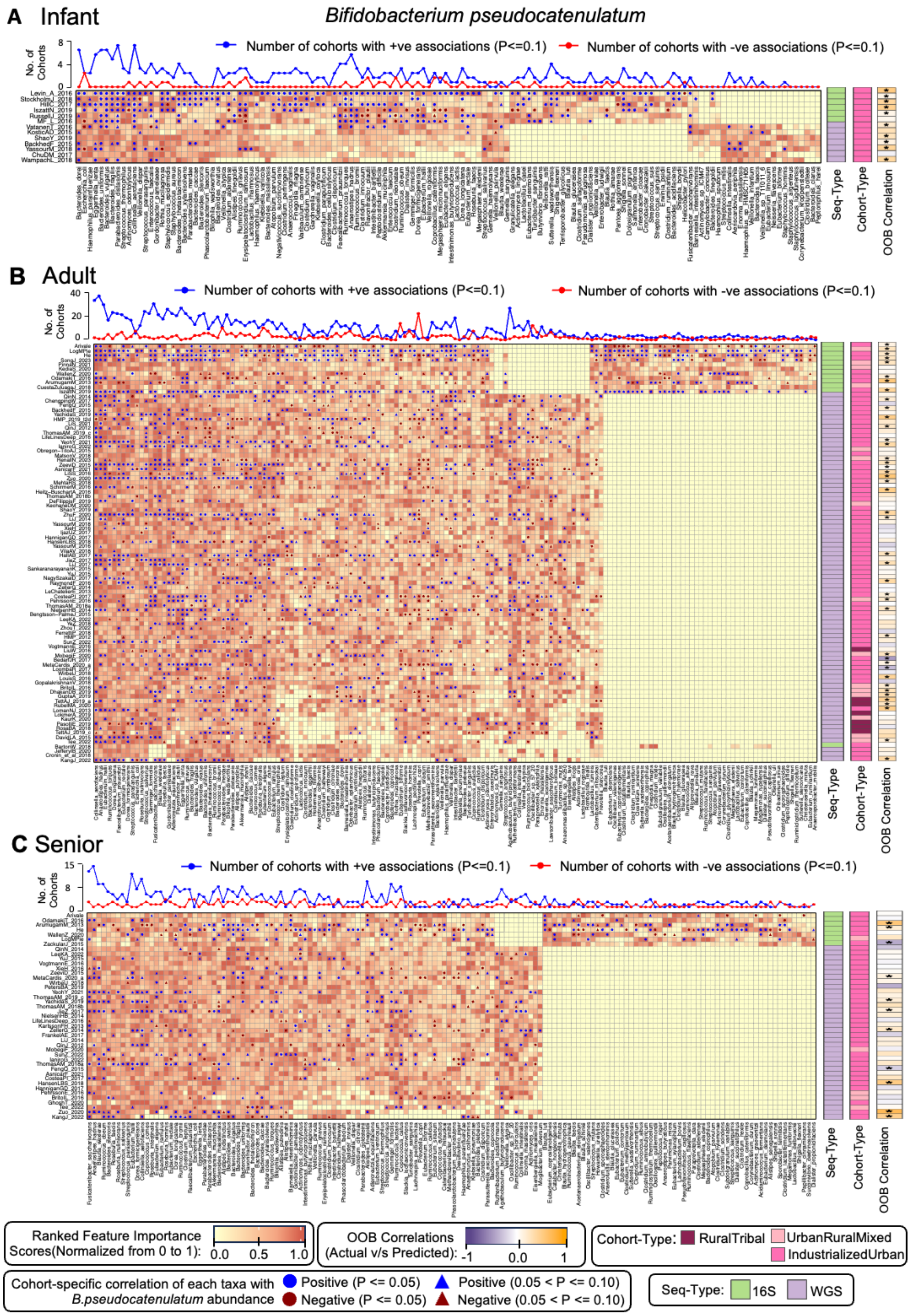


**Figure S6. Random Forest (RF) model-based identification of distinct sets of non-Bifidobacterial taxa that consistently associated with the abundance of *Bifidobacterium pseudocatenulatum*, serving as top predictors of *B. pseudocatenulatum* abundance in the gut microbiome. A-C.** Heatmap showing the variation of the rank-normalized feature importance scores of the top predictor taxa for the Random Forest models built for Infant (A), Adult (B) and Senior (C) cohorts. The profiling strategy (Seq-Type), and the life-style (Cohort-Type) of each cohort is indicated as strips as shown for each heatmap. Also indicated are the out-of-bag (OOB) correlations between the actual and RF-predicted *B. pseudocatenulatum* for each cohort. Symbols within the heatmap cells indicate both the direction (positive or negative) and strength (based on p-values) of the association between each non-Bifidobacterial taxon (columns) and *B. pseudocatenulatum* abundance within each cohort (rows). Line plots above each heatmap summarize these associations and show the number of cohorts where each non-Bifidobacterial taxon displays significant positive or negative associations with *B. pseudocatenulatum* abundance ($p \leq 0.1$). The extent and consistency of these associations varied by the specific non-Bifidobacterial taxon, with some showing notably higher predictive power and consistent directionality. Association patterns were also cohort-type-dependent, namely Infant cohorts exhibited distinct sets of associated taxa compared to other age groups (same for other age-groups); 16S and WGS cohorts showed differing patterns. We also observed variations depending upon the life-style of cohorts. These patterns were captured in the Association-Scores as described in **Figure 3**.

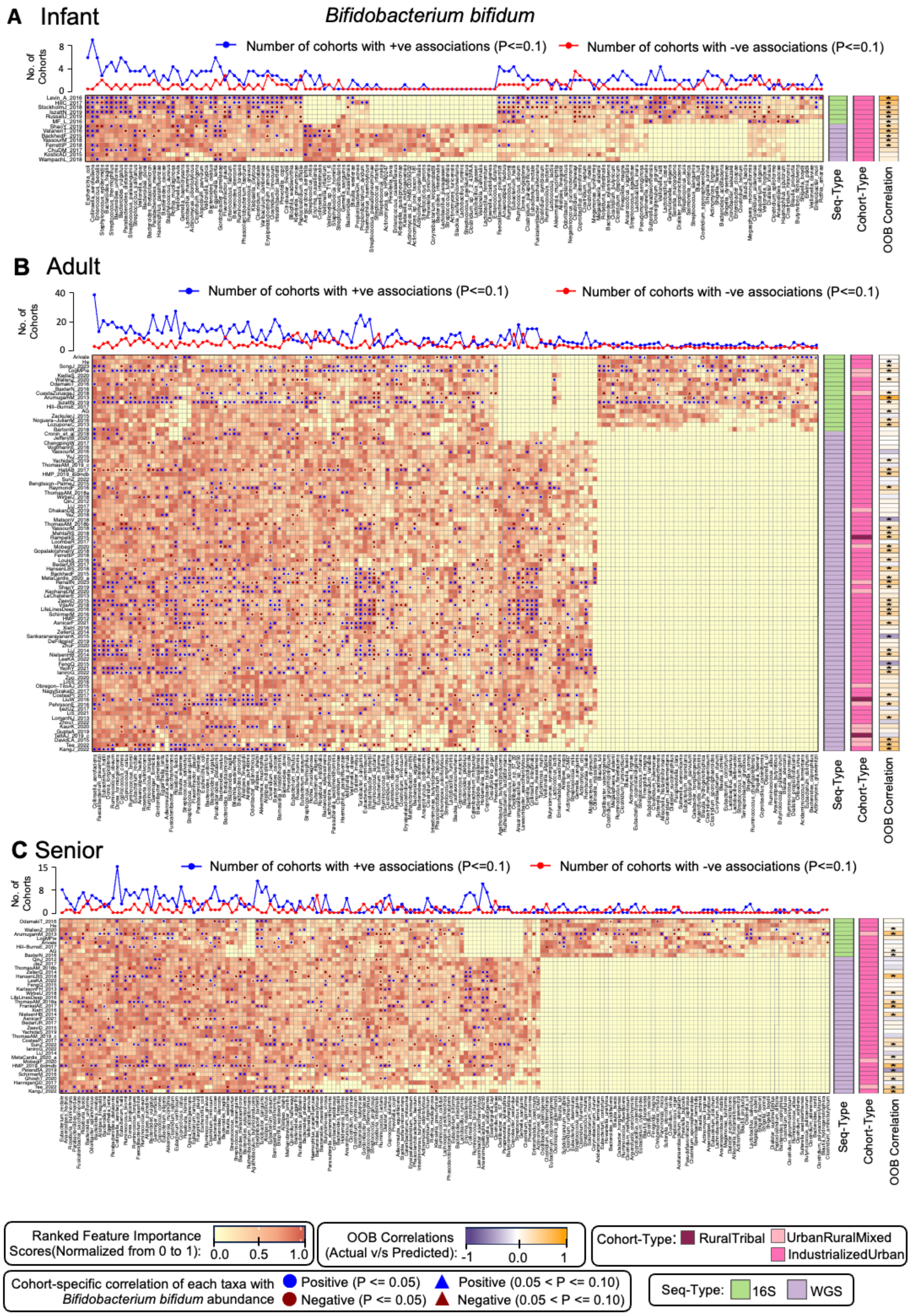


**Figure S7. Random Forest (RF) model-based identification of distinct sets of non-Bifidobacterial taxa that consistently associated with the abundance of *Bifidobacterium bifidum*, serving as top predictors of *B. bifidum* abundance in the gut microbiome. A-C.** Heatmap showing the variation of the rank-normalized feature importance scores of the top predictor taxa for the Random Forest models built for Infant (A), Adult (B) and Senior (C) cohorts. The profiling strategy (Seq-Type), and the life-style (Cohort-Type) of each cohort is indicated as strips as shown for each heatmap. Also indicated are the out-of-bag (OOB) correlations between the actual and RF-predicted *B. bifidum* for each cohort. Symbols within the heatmap cells indicate both the direction (positive or negative) and strength (based on p-values) of the association between each non-Bifidobacterial taxon (columns) and *B. bifidum* abundance within each cohort (rows). Line plots above each heatmap summarize these associations and show the number of cohorts where each non-Bifidobacterial taxon displays significant positive or negative associations with *B. bifidum* abundance ($p \leq 0.1$). The extent and consistency of these associations varied by the specific non-Bifidobacterial taxon, with some showing notably higher predictive power and consistent directionality. Association patterns were also cohort-type-dependent, namely Infant cohorts exhibited distinct sets of associated taxa compared to other age groups (same for other age-groups); 16S and WGS cohorts showed differing patterns. We also observed variations depending upon the life-style of cohorts. These patterns were captured in the Association-Scores as described in **Figure 3**.

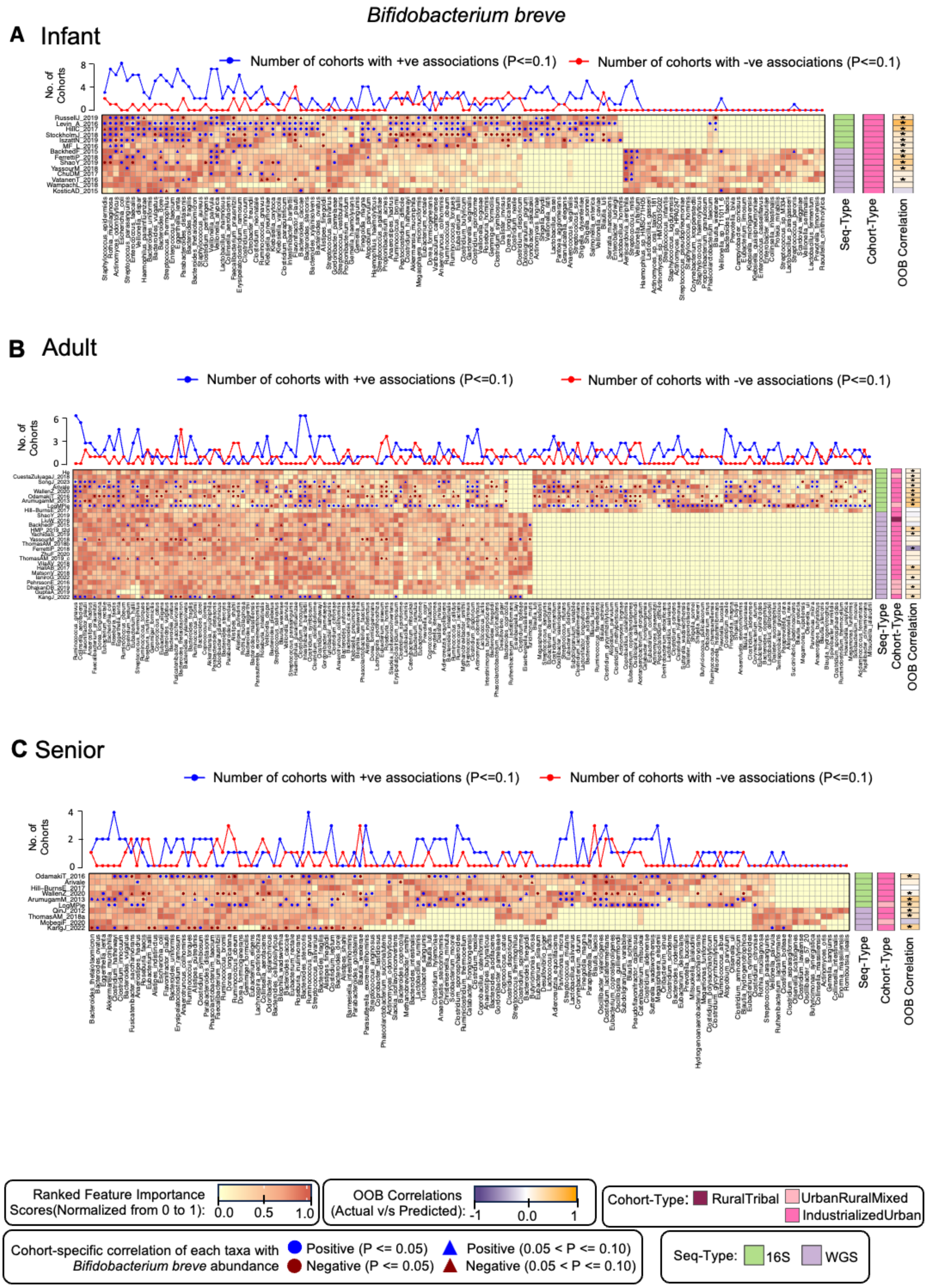


**Figure S8. Random Forest (RF) model-based identification of distinct sets of non-Bifidobacterial taxa that consistently associated with the abundance of *Bifidobacterium breve*, serving as top predictors of *B. breve* abundance in the gut microbiome. A-C.**

Heatmap showing the variation of the rank-normalized feature importance scores of the top predictor taxa for the Random Forest models built for Infant (A), Adult (B) and Senior (C) cohorts. The profiling strategy (Seq-Type), and the life-style (Cohort-Type) of each cohort is indicated as strips as shown for each heatmap. Also indicated are the out-of-bag (OOB) correlations between the actual and RF-predicted *B. breve* for each cohort. Symbols within the heatmap cells indicate both the direction (positive or negative) and strength (based on p-values) of the association between each non-Bifidobacterial taxon (columns) and *B. breve* abundance within each cohort (rows). Line plots above each heatmap summarize these associations and show the number of cohorts where each non-Bifidobacterial taxon displays significant positive or negative associations with *B. breve* abundance ($p \leq 0.1$). The extent and consistency of these associations varied by the specific non-Bifidobacterial taxon, with some showing notably higher predictive power and consistent directionality. Association patterns were also cohort-type-dependent, namely Infant cohorts exhibited distinct sets of associated taxa compared to other age groups (same for other age-groups); 16S and WGS cohorts showed differing patterns. We also observed variations depending upon the life-style of cohorts. These patterns were captured in the Association-Scores as described in **Figure 3**.

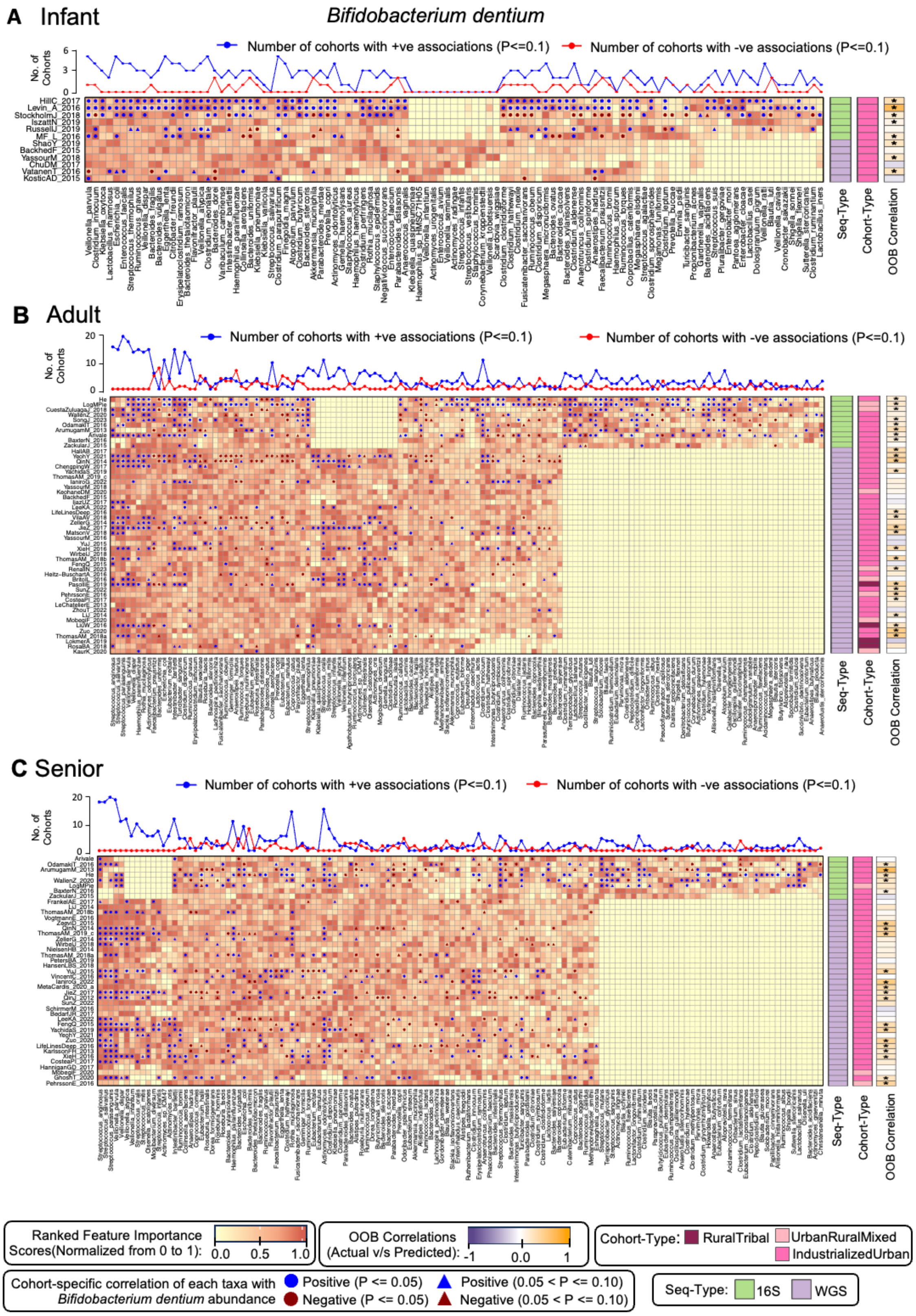


**Figure S9. Random Forest (RF) model-based identification of distinct sets of non-Bifidobacterial taxa that consistently associated with the abundance of *Bifidobacterium dentium*, serving as top predictors of *B. dentium* abundance in the gut microbiome.** A-C. Heatmap showing the variation of the rank-normalized feature importance scores of the top predictor taxa for the Random Forest models built for Infant (A), Adult (B) and Senior (C) cohorts. The profiling strategy (Seq-Type), and the life-style (Cohort-Type) of each cohort is indicated as strips as shown for each heatmap. Also indicated are the out-of-bag (OOB) correlations between the actual and RF-predicted *B. dentium* abundance for each cohort. Symbols within the heatmap cells indicate both the direction (positive or negative) and strength (based on p-values) of the association between each non-Bifidobacterial taxon (columns) and *B. dentium* abundance within each cohort (rows). Line plots above each heatmap summarize these associations and show the number of cohorts where each non-Bifidobacterial taxon displays significant positive or negative associations with *B. dentium* abundance ($p \leq 0.1$). The extent and consistency of these associations varied by the specific non-Bifidobacterial taxon, with some showing notably higher predictive power and consistent directionality. Association patterns were also cohort-type-dependent, namely Infant cohorts exhibited distinct sets of associated taxa compared to other age groups (same for other age-groups); 16S and WGS cohorts showed differing patterns. We also observed variations depending upon the life-style of cohorts. These patterns were captured in the Association-Scores as described in **Figure 3**.

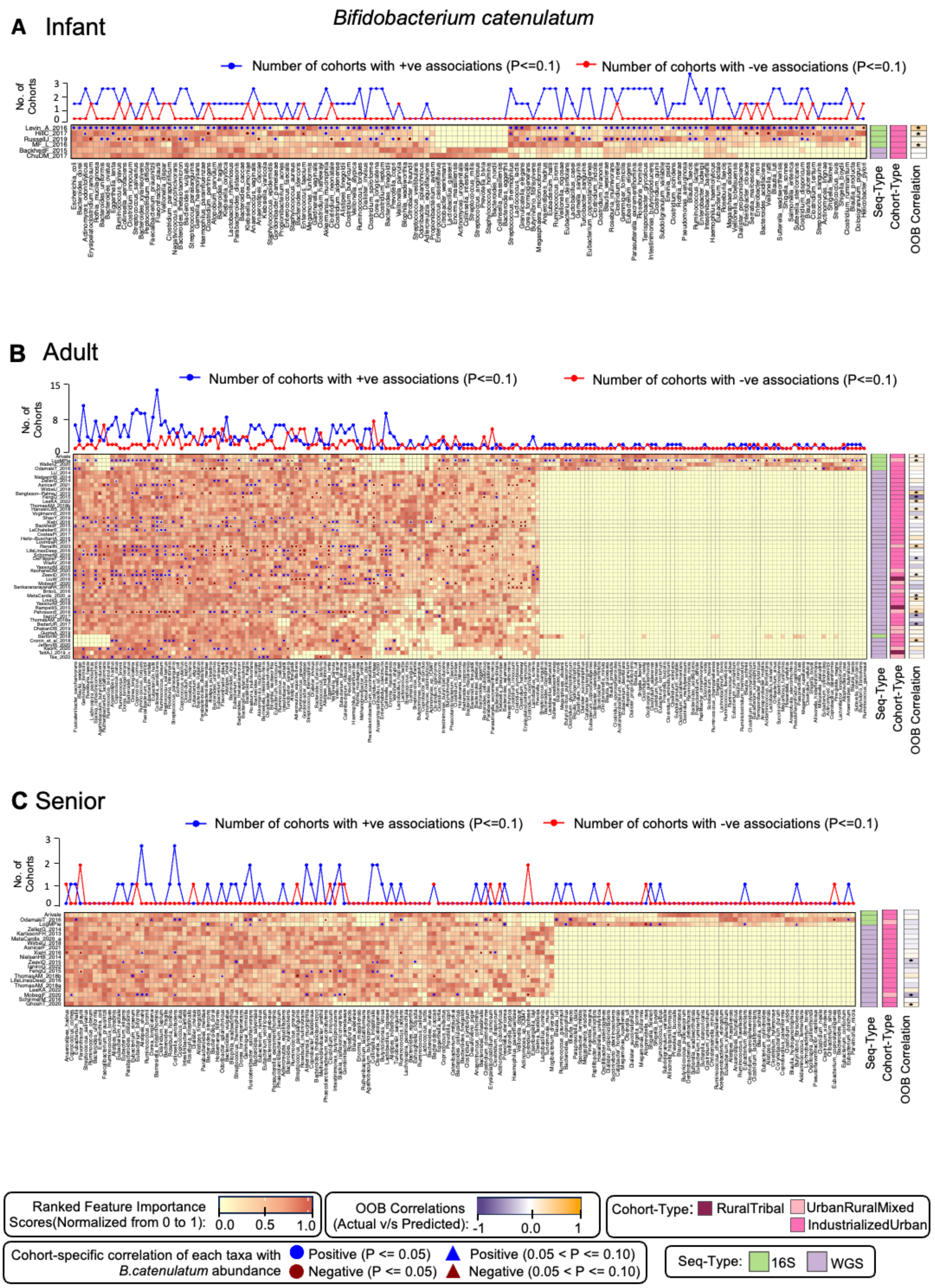


**Figure S10. Random Forest (RF) model-based identification of distinct sets of non-Bifidobacterial taxa that consistently associated with the abundance of *Bifidobacterium catenulatum*, serving as top predictors of *B. catenulatum* abundance in the gut microbiome. A-C.** Heatmap showing the variation of the rank-normalized feature importance scores of the top predictor taxa for the Random Forest models built for Infant (A), Adult (B)

and Senior (C) cohorts. The profiling strategy (Seq-Type), and the life-style (Cohort-Type) of each cohort is indicated as strips as shown for each heatmap. Also indicated are the out-of-bag (OOB) correlations between the actual and RF-predicted *B. catenulatum* for each cohort. Symbols within the heatmap cells indicate both the direction (positive or negative) and strength (based on p-values) of the association between each non-Bifidobacterial taxon (columns) and *B. catenulatum* abundance within each cohort (rows). Line plots above each heatmap summarize these associations and show the number of cohorts where each non-Bifidobacterial taxon displays significant positive or negative associations with *B. catenulatum* abundance ($p \leq 0.1$). The extent and consistency of these associations varied by the specific non-Bifidobacterial taxon, with some showing notably higher predictive power and consistent directionality. Association patterns were also cohort-type-dependent, namely Infant cohorts exhibited distinct sets of associated taxa compared to other age groups (same for other age-groups); 16S and WGS cohorts showed differing patterns. We also observed variations depending upon the life-style of cohorts. These patterns were captured in the Association-Scores as described in **Figure 3**.

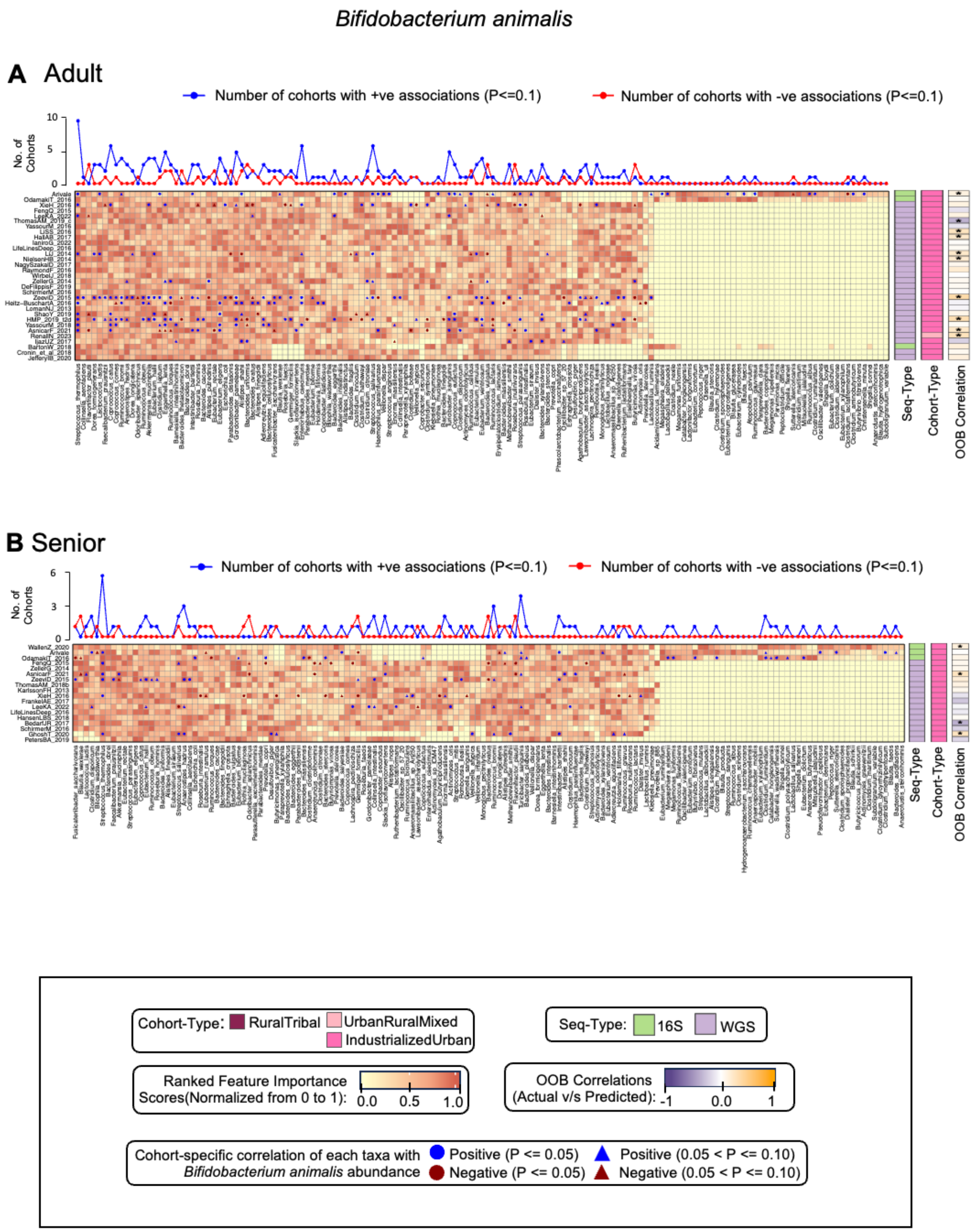


**Figure S11. Random Forest (RF) model-based identification of distinct sets of non-Bifidobacterial taxa that consistently associated with the abundance of *Bifidobacterium animalis*, serving as top predictors of *B. animalis* abundance in the gut microbiome. A-B.** Heatmap showing the variation of the rank-normalized feature importance scores of the top predictor taxa for the Random Forest models built for Adult (A) and Senior (B) cohorts. The profiling strategy (Seq-Type), and the life-style (Cohort-Type) of each cohort is indicated as strips as shown for each heatmap. Also indicated are the out-of-bag (OOB) correlations between the actual and RF-predicted *B. animalis* abundance for each cohort. Symbols within the heatmap cells indicate both the direction (positive or negative) and strength (based on p-values) of the association between each non-Bifidobacterial taxon (columns) and *B. animalis* abundance within each cohort (rows). Line plots above each heatmap summarize these associations and show the number of cohorts where each non-Bifidobacterial taxon displays significant positive or negative associations with *B. animalis* abundance ($p \leq 0.1$). The extent and consistency of these associations varied by the specific non-Bifidobacterial taxon, with some showing notably higher predictive power and consistent directionality. Association patterns were also cohort-type-dependent, namely Infant cohorts exhibited distinct sets of associated taxa compared to other age groups (same for other age-groups); 16S and WGS cohorts showed differing patterns. We also observed variations depending upon the life-style of cohorts. These patterns were captured in the Association-Scores as described in **Figure 3**.

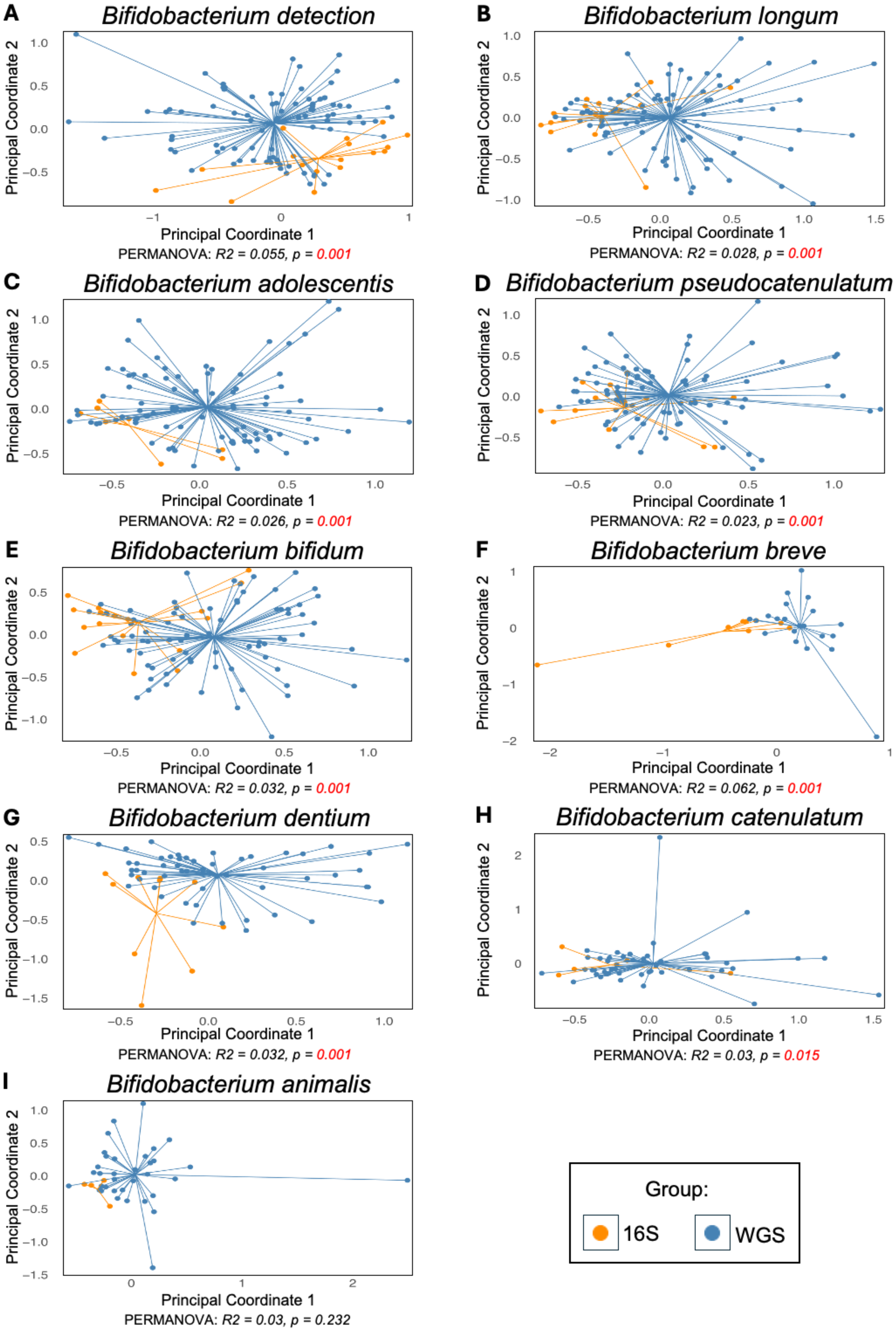

A
Bifidobacterium detection
Principal Coordinate 2
Principal Coordinate 1
PERMANOVA: R2 = 0.055, p = 0.001
B
Bifidobacterium longum
Principal Coordinate 2
Principal Coordinate 1
PERMANOVA: R2 = 0.028, p = 0.001
C
Bifidobacterium adolescentis
Principal Coordinate 2
Principal Coordinate 1
PERMANOVA: R2 = 0.026, p = 0.001
D
Bifidobacterium pseudocatenulatum
Principal Coordinate 2
Principal Coordinate 1
PERMANOVA: R2 = 0.023, p = 0.001
E
Bifidobacterium bifidum
Principal Coordinate 2
Principal Coordinate 1
PERMANOVA: R2 = 0.032, p = 0.001
F
Bifidobacterium breve
Principal Coordinate 2
Principal Coordinate 1
PERMANOVA: R2 = 0.062, p = 0.001
G
Bifidobacterium dentium
Principal Coordinate 2
Principal Coordinate 1
PERMANOVA: R2 = 0.032, p = 0.001
H
Bifidobacterium catenulatum
Principal Coordinate 2
Principal Coordinate 1
PERMANOVA: R2 = 0.03, p = 0.015
I
Bifidobacterium animalis
Principal Coordinate 2
Principal Coordinate 1
PERMANOVA: R2 = 0.03, p = 0.232
Group:
16S
WGS

**Figure S12. Principal Coordinate Analysis (PCoA) results showing the variation of the ranked Feature Importance Vectors for the cohort-specific Random Forest models for the different cohorts based on their profiling strategy (Seq-Type; i.e. 16S or WGS).** The Feature Importance vectors contain the ranked Feature Importance scores for the different non-Bifidobacterial taxa obtained for the Random Forest model generated for that cohort for the prediction of **A.** Overall *Bifidobacterium* detection **B.** *B. longum* **C.** *B. adolescentis* **D.** *B. pseudocatenulatum* **E.** *B. bifidum* **F.** *B. breve* **G.** *B. dentium* and **H.** *B. catenulatum* **I.** *B. animalis*, using adult/senior studies. 16S and WGS cohorts are denoted as orange and blue-points, respectively and connected with lines to their group centroid.

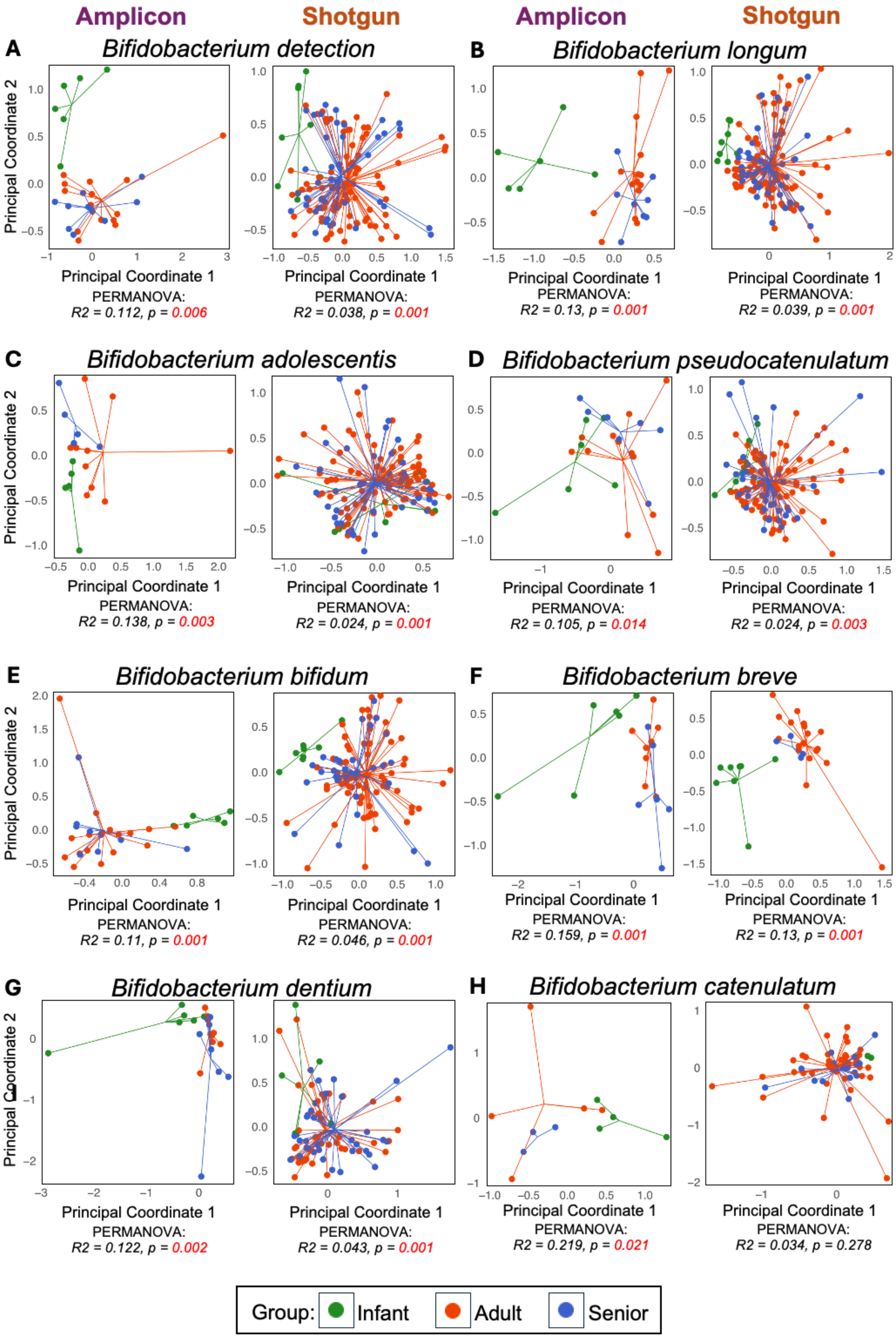

Amplicon
Shotgun
Amplicon
Shotgun
A
Bifidobacterium detection
PERMANOVA:
R2 = 0.112, p = 0.006
PERMANOVA:
R2 = 0.038, p = 0.001
B
Bifidobacterium longum
PERMANOVA:
R2 = 0.13, p = 0.001
PERMANOVA:
R2 = 0.039, p = 0.001
C
Bifidobacterium adolescentis
PERMANOVA:
R2 = 0.138, p = 0.003
PERMANOVA:
R2 = 0.024, p = 0.001
D
Bifidobacterium pseudocatenulatum
PERMANOVA:
R2 = 0.105, p = 0.014
PERMANOVA:
R2 = 0.024, p = 0.003
E
Bifidobacterium bifidum
PERMANOVA:
R2 = 0.11, p = 0.001
PERMANOVA:
R2 = 0.046, p = 0.001
F
Bifidobacterium breve
PERMANOVA:
R2 = 0.159, p = 0.001
PERMANOVA:
R2 = 0.13, p = 0.001
G
Bifidobacterium dentium
PERMANOVA:
R2 = 0.122, p = 0.002
PERMANOVA:
R2 = 0.043, p = 0.001
H
Bifidobacterium catenulatum
PERMANOVA:
R2 = 0.219, p = 0.021
PERMANOVA:
R2 = 0.034, p = 0.278
Principal Coordinate 1
Principal Coordinate 2
Group: Infant Adult Senior

**Figure S13. Principal Coordinate Analysis (PCoA) results showing the variation of the ranked Feature Importance Vectors for the cohort-specific Random Forest models for the different cohorts based on their age-group (i.e. Infants, Adults and Seniors).** The Feature Importance vectors contain the ranked Feature Importance scores for the different non-Bifidobacterial taxa obtained for the Random Forest model generated for that cohort for the prediction of **A.** Overall *Bifidobacterium* detection **B.** *B. longum* **C.** *B. adolescentis* **D.** *B. pseudocatenulatum* **E.** *B. bifidum* **F.** *B. breve* **G.** *B. dentium* and **H.** *B. catenulatum*. Each point refers to a study-cohort, with the infant, adult and senior indicated in green, red and blue colors, respectively, and connected by lines to the corresponding group centroids. PERMANOVA results (shown using both the R2 and p-value) shows statistically significant separation for all species except *B. catenulatum,* in WGS data.

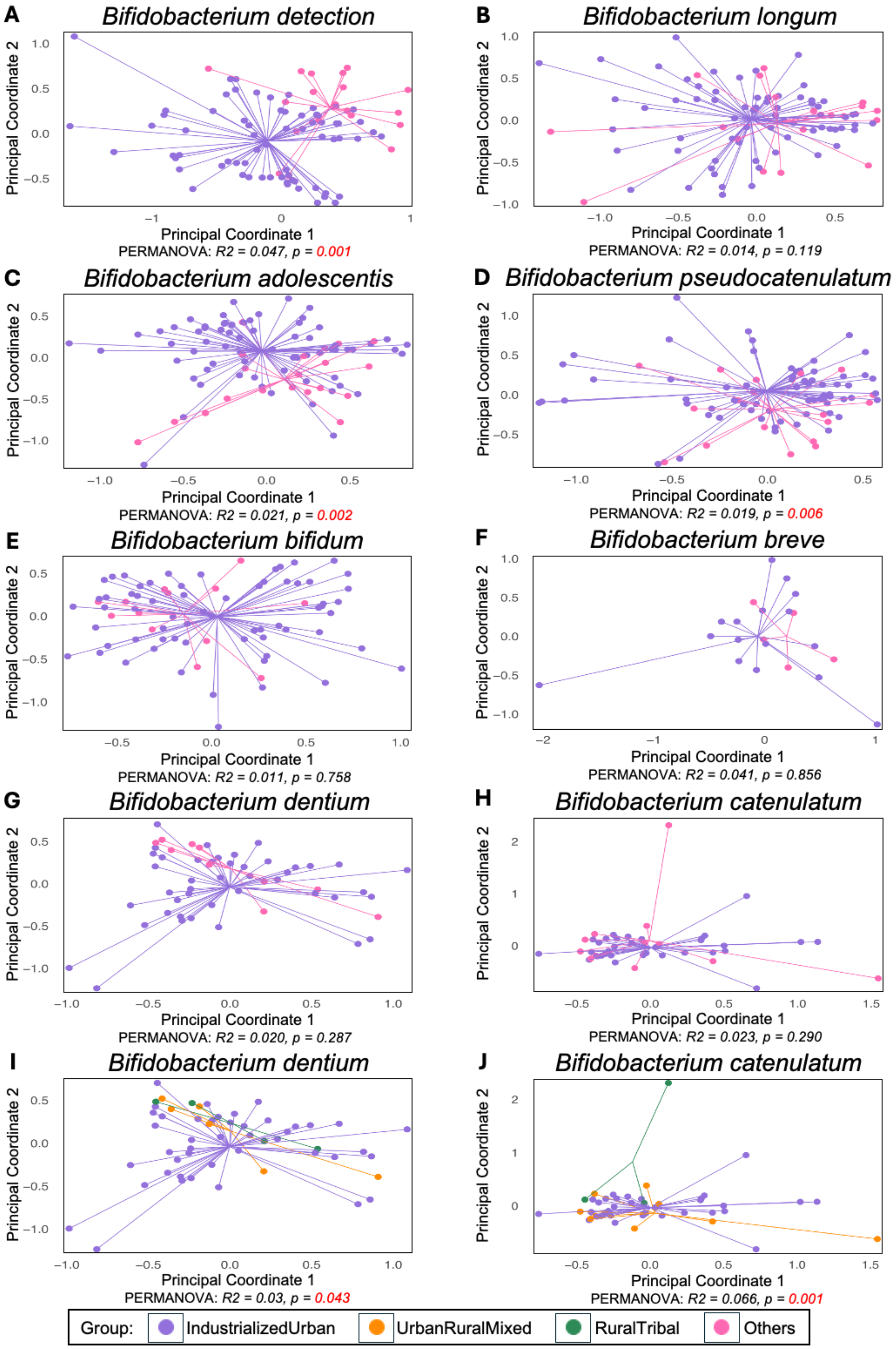
A
Bifidobacterium detection
Principal Coordinate 2
Principal Coordinate 1
PERMANOVA: R2 = 0.047, p = 0.001
B
Bifidobacterium longum
Principal Coordinate 2
Principal Coordinate 1
PERMANOVA: R2 = 0.014, p = 0.119
C
Bifidobacterium adolescentis
Principal Coordinate 2
Principal Coordinate 1
PERMANOVA: R2 = 0.021, p = 0.002
D
Bifidobacterium pseudocatenulatum
Principal Coordinate 2
Principal Coordinate 1
PERMANOVA: R2 = 0.019, p = 0.006
E
Bifidobacterium bifidum
Principal Coordinate 2
Principal Coordinate 1
PERMANOVA: R2 = 0.011, p = 0.758
F
Bifidobacterium breve
Principal Coordinate 2
Principal Coordinate 1
PERMANOVA: R2 = 0.041, p = 0.856
G
Bifidobacterium dentium
Principal Coordinate 2
Principal Coordinate 1
PERMANOVA: R2 = 0.020, p = 0.287
H
Bifidobacterium catenulatum
Principal Coordinate 2
Principal Coordinate 1
PERMANOVA: R2 = 0.023, p = 0.290
I
Bifidobacterium dentium
Principal Coordinate 2
Principal Coordinate 1
PERMANOVA: R2 = 0.03, p = 0.043
J
Bifidobacterium catenulatum
Principal Coordinate 2
Principal Coordinate 1
PERMANOVA: R2 = 0.066, p = 0.001
Group: IndustrializedUrban UrbanRuralMixed RuralTribal Others

**Figure S14. Principal Coordinate Analysis (PCoA) results showing the variation of the ranked Feature Importance Vectors for the cohort-specific Random Forest models for the different cohorts based on their Lifestyle-associated Cohort Type**. This analysis was restricted to WGS adult and senior cohorts due to the absence of Non-IndustrializedUrban cohorts (i.e., UrbanRuralMixed and RuralTribal) in the infant age group and their markedly lower representation in the 16S datasets. Feature Importance vectors contain the ranked Feature Importance scores for the different non-Bifidobacterial taxa obtained for the Random Forest model generated for that cohort for the prediction of **A.** Overall *Bifidobacterium* detection **B.** *B. longum* **C.** *B. adolescentis* **D.** *B. pseudocatenulatum* **E.** *B. bifidum* **F.** *B. breve* **G.** *B. dentium* **H.** *B. catenulatum.* In panels **A–H**, each point represents a study-cohort, coloured by group: IndustrializedUrban cohorts in purple and Non-IndustrializedUrban cohorts (combining RuralTribal and UrbanRuralMixed) in pink. Panels **I** and **J** denote the same results for B. *dentium* and B. *catenulatum*, respectively, where in the Other Non-IndustrializedUrban cohorts are resolved into UrbanRuralMixed (orange) and RuralTribal (green). PERMANOVA results are reported with $R^2$ (strength) and p-values (significance). Significant separation of feature importance vectors was observed in between the IndustrializedUrban and Other Non-Industrialized cohorts for overall *Bifidobacterium* detection, *B. adolescentis*, *B. pseudocatenulatum*. When UrbanRuralMixed and RuralTribal were separated, significant separation was also observed for *B. dentium* and *B. catenulatum* indicating five of the nine Bifidobacterial traits showed significant variations of the association patterns with non-Bifidobacterial taxa, depending upon the life-style of the investigated cohort.

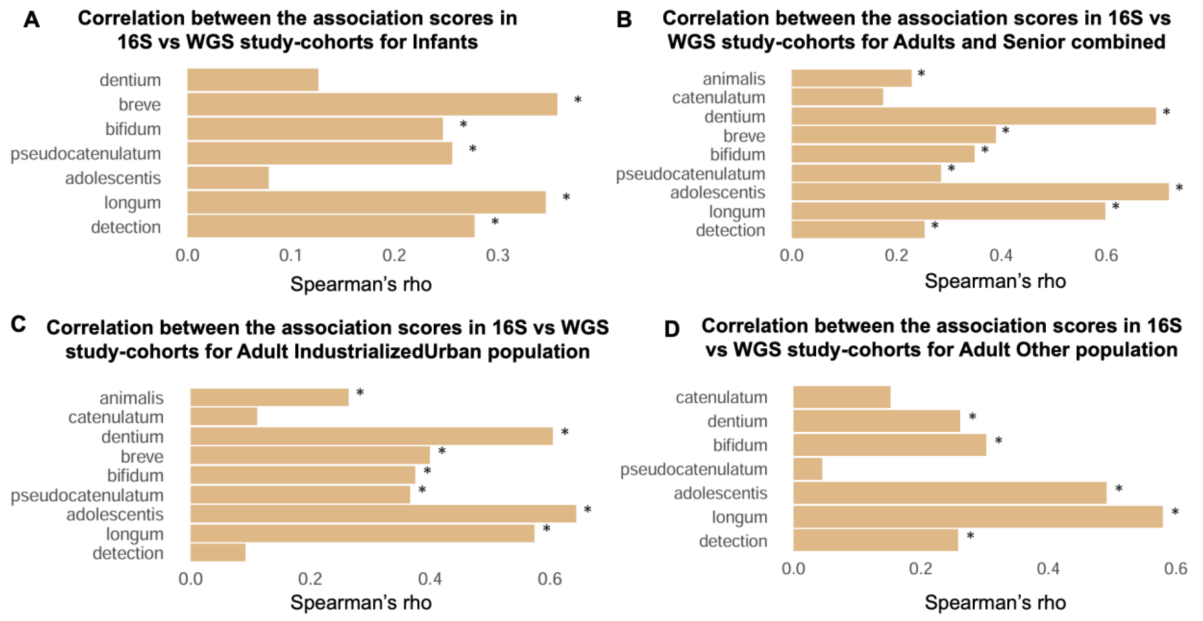


**Figure S15.** Correlation between Association-Scores (obtained for different non-Bifidobacterial taxa with respect to various Bifidobacterial traits) derived from 16S and WGS data in (A) Infant cohort-groups (B) Adult & Senior cohort-groups (C) IndustrializedUrban cohort groups and (D) Other Non IndustrializedUrban cohort groups (UrbanRuralMixed + RuralTribal). Associations among overall *Bifidobacterium* detection and different Bifidobacterial taxa using the Association-Scores with non-bifidobacterial taxa in 16S vs. WGS study-cohorts. Spearman's correlation coefficient has been shown in the form of bar plots for overall Bifidobacterial taxa detection and different Bifidobacterial taxa individually between 16S and WGS profiling-strategy-based cohorts having microbiomes from infants (A), adults and senior combined (B), microbiomes from IndustrializedUrban population (C) and in other populations including UrbanRuralMixed and RuralTribal combined (D) using the Association-Scores with non-bifidobacterial taxa. The statistically significant ($p<=0.05$) associations have been marked as '*' on each bar. Some Bifidobacteria (*B. catenulatum* and *B. animalis* in infants; *B. breve* and *B. animalis* in other Non-IndustrializedUrban cohorts) don't show consistent association pattern with the non-Bifidobacterial taxa across either profiling strategy, hence, they have not been included.

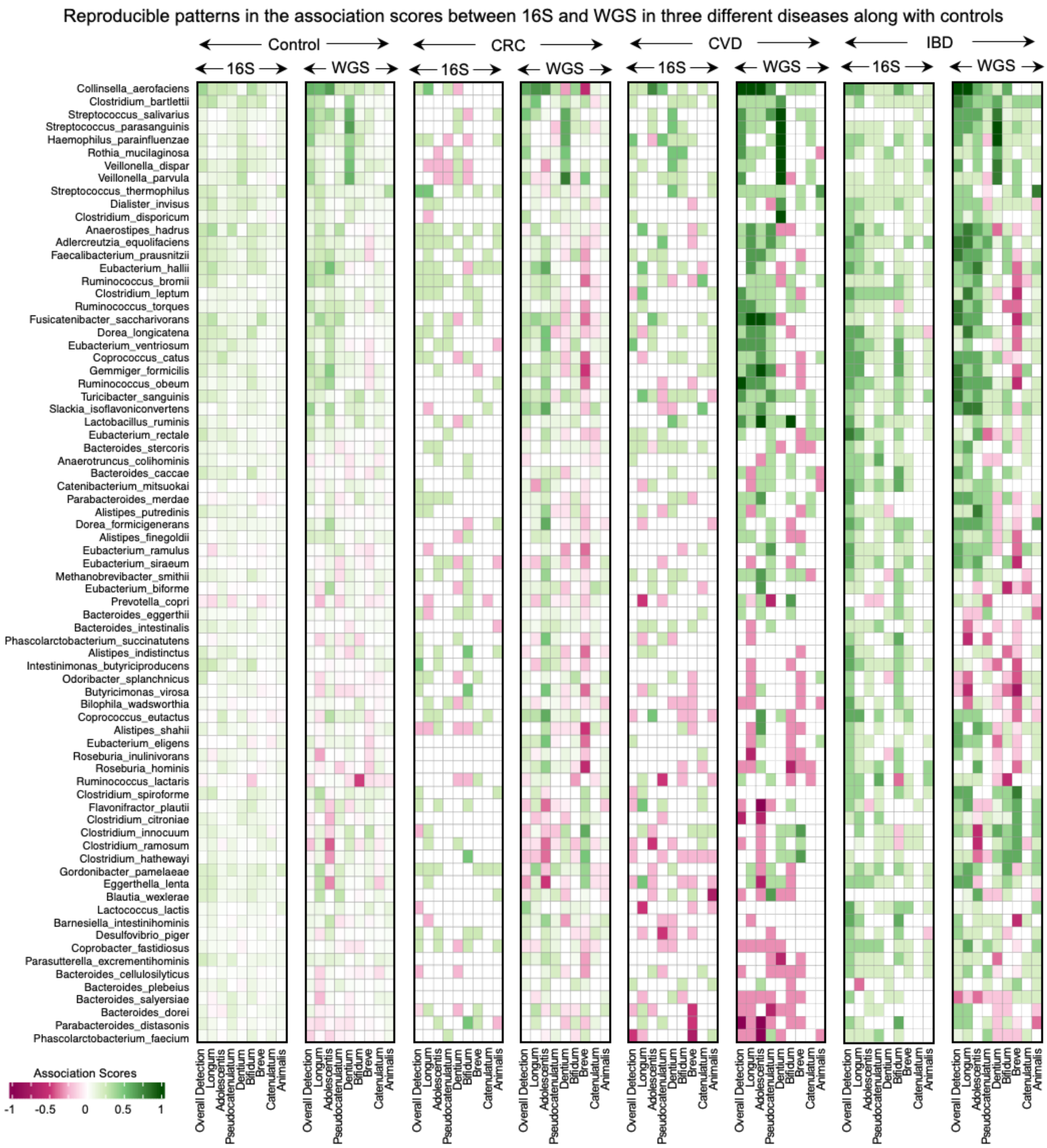


**Figure S16. Association patterns of the non-Bifidobacterial taxa with the different Bifidobacterial properties show reasonable similar patterns across study-cohorts from different profiling strategy (16S and WGS) in majority of the Bifidobacterium-trait-Disease combinations.** Heatmap showing the variation of Association-Score patterns of the non-Bifidobacterial taxa with different Bifidobacterial traits computed separately for 16S and WGS study-cohorts corresponding to the non-diseased controls (matched controls from each of the study-cohorts) and the four diseased groups (as shown). Only those diseases were considered where data was available from more than three cohorts in each of the sequencing type category (16S and WGS) to investigate the reproducibility of the Association-Score across both the profiling strategy. Only those non-Bifidobacterial taxa showing consistent association patterns (Association-Score >= 0.25) with the different Bifidobacterial traits in more than two Bifidobacterium-trait-Disease combinations are shown here. The colour scale indicates the range of Association-Scores spanning from -1 to 1.

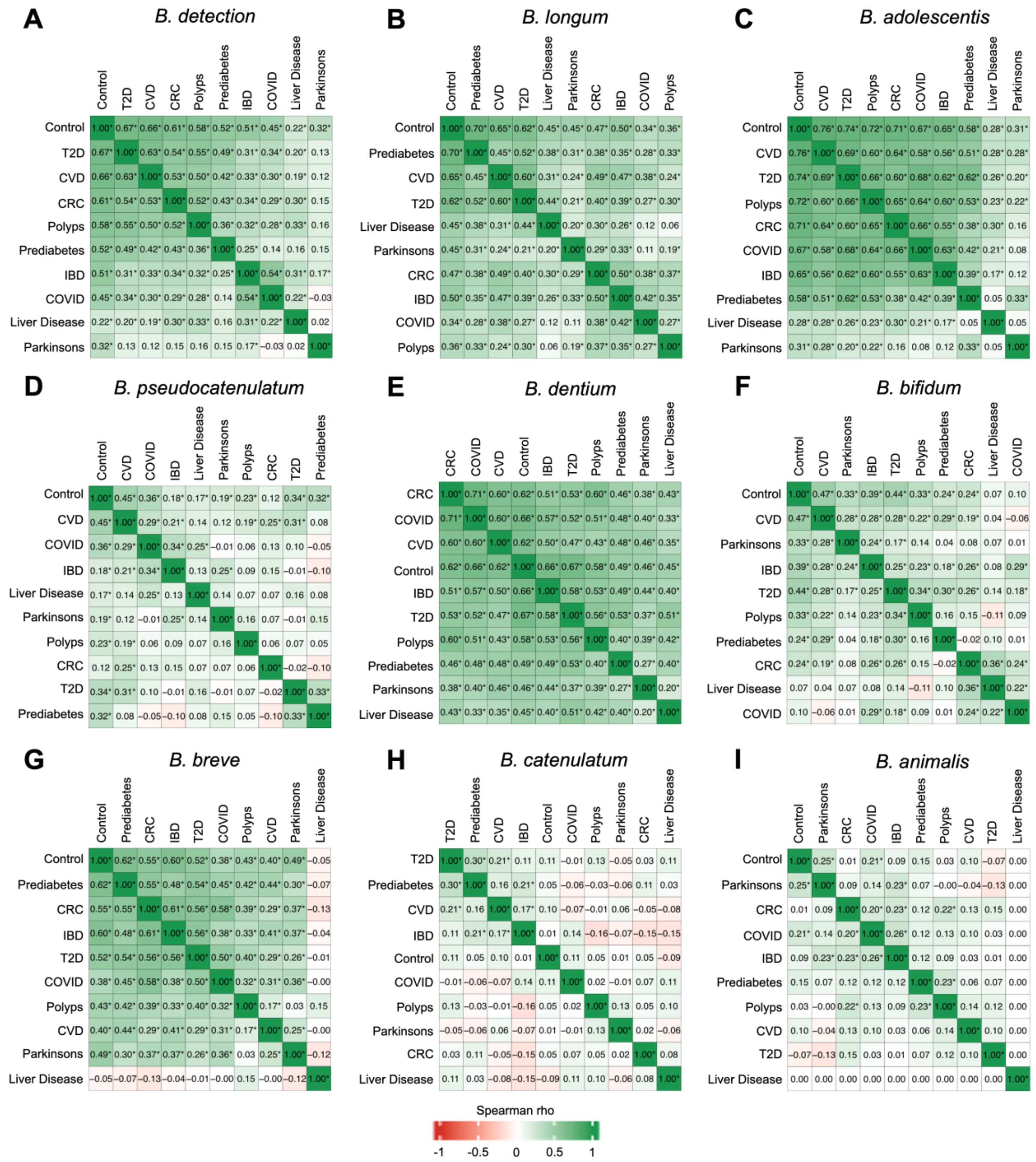


**Figure S17.** Correlations between the Association-Scores obtained for the different non-Bifidobacterial taxa in the study-sub-cohorts for the nine diseases and the matched-controls with respect to the nine Bifidobacterium properties, namely **A.** *Bifidobacterium* detection **B.** *B. longum* **C.** *B. adolescentis* **D.** *B. pseudocatenulatum* **E.** *B. dentium* **F.** *B. bifidum* **G.** *B. breve* **H.** *B. catenulatum* **I.** *B. animalis*. Association-Scores of non-Bifidobacterium taxa with *Bifidobacterium* properties were computed separately for apparently non-diseased sub-cohorts (disease-matched controls) and in the study-sub-cohorts containing microbiomes from nine diseases. For each *Bifidobacterium* property, for each pair of groups (10 conditions in total: 9 diseases + control), the spearman correlation coefficients were calculated between the Association-Scores obtained for the different non-Bifidobacterium taxa. The Heatmaps displays these pairwise correlation values, with positive correlations in green and negative correlations in red, scaled between –1 and 1. * denote statistically significant correlations ($p \leq 0.05$). Distinct patterns emerge between control and disease states as well as across different diseases, highlighting disease-specific alterations in *Bifidobacterium* taxa association landscapes.

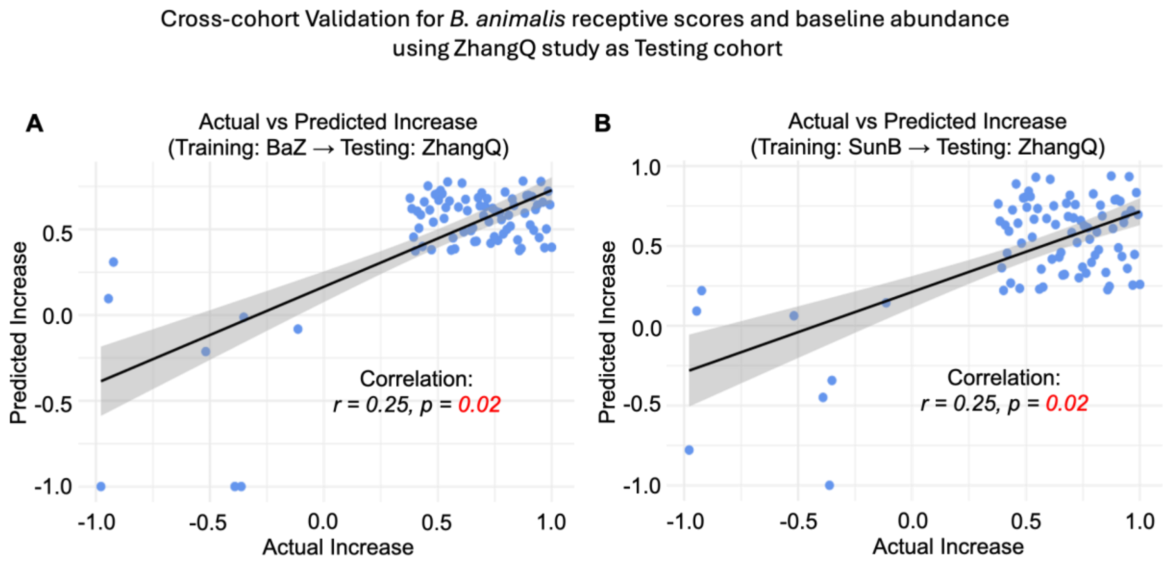


**Figure S18. Cross-cohort validation of *Bifidobacterium animalis* receptive scores and baseline abundance models using ZhangQ as the testing cohort.** Validation of predictive models for post-treatment increase in *B. animalis* using receptive scores and baseline abundance derived from independent cohorts. **A.** Model trained on BaZ study **B.** Model trained on SunB study. In both cases, ZhangQ cohort was used for testing. Spearman correlation coefficients (rho) and corresponding *p*-values indicate the strength and significance of the association between predicted and observed *B. animalis* increase across trials.

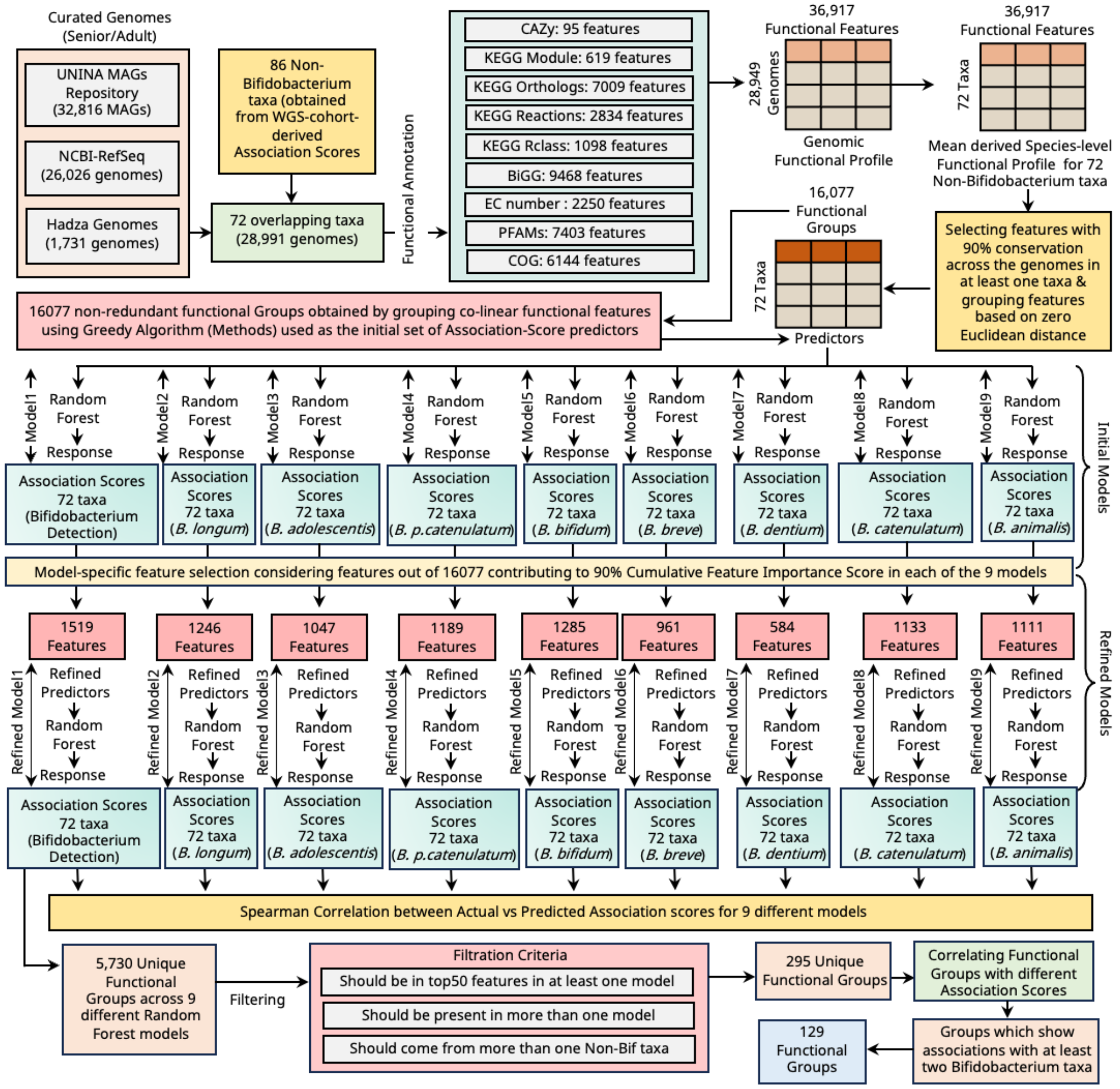


**Figure S19.** This panel shows the schematic workflow of how the mean functional feature abundance profile is generated using a set of 28,991 high quality genomes including MAGs generated from Senior/Adults microbiomes from different genomic repositories, then it shows the approach of how the co-linear functional features were grouped into unique functional-groups using a Greedy algorithm. Furthermore, the random forest-based feature selection and identifying top features which constitute 90% cumulative feature importance scores in each of the nine models (eight Bifidobacterial taxa and one total detection), followed by re-training of the random forest-based models with the top features to reduce noise and finally correlating the actual and predicted Association-Scores using Spearman correlation. The unique top functional-groups coming from nine different models were further reduced based on three selection criteria and then associating the most relevant features to each of the nine types of Association-Scores to capture the association patterns.

Association Patterns of Non-Bifidobacterium Taxa derived Functions with the association scores of total Bifidobacterial Taxa detection and eight different Bifidobacteria

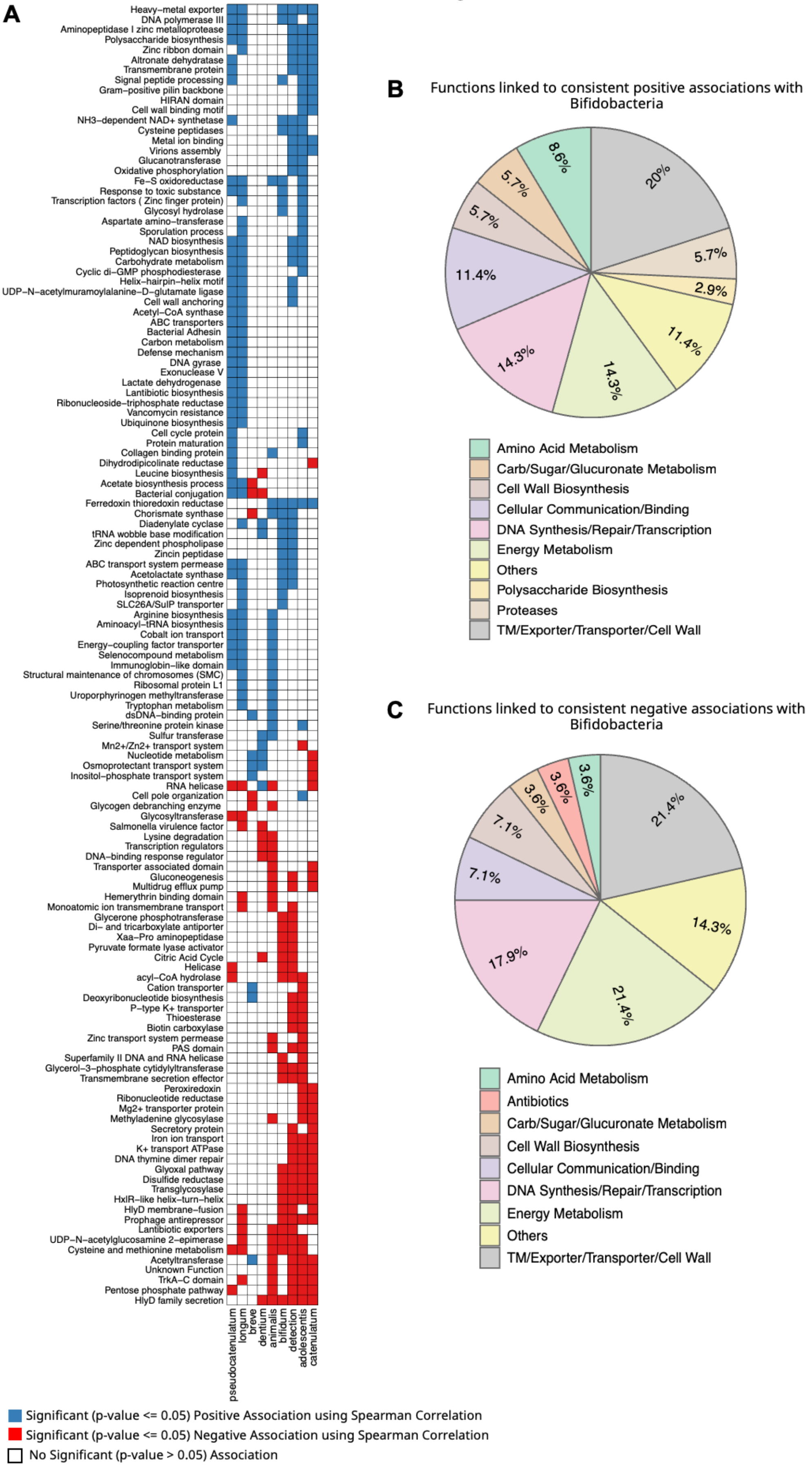

**Figure S20. Association patterns of non-Bifidobacterial taxa derived functions with eight different Bifidobacterial taxa and total Bifidobacterial detection. A.** The heatmap shows the associations-patterns where blue colour cells indicate significant (p-value <= 0.05) positive association (Spearman correlation) and on the other hand red coloured cells indicate significant (p-value <= 0.05) negative association and white cells show no statistically significant (p-value > 0.05) associations. **B.** The pie-chart shows the relative percentages of the functions linked to consistent positive associations with multiple Bifidobacterial properties **C.** The relative percentages of the functions linked to consistent negative associations with multiple Bifidobacterial properties have been shown in this pie-chart.
